# Southern (California) Sea Otter Population Status and Trends at San Nicolas Island, 2023–Winter 2026

**DOI:** 10.64898/2026.08.28.747881

**Authors:** By Joseph A. Tomoleoni, Julie L. Yee, Elizabeth Seacord, Michelle M. Staedler, Brian B. Hatfield, Lilian Carswell, Jessica Fujii, Gena B. Bentall, Leilani Konrad, Colleen Young, M. Tim Tinker, Lizabeth Bowen

## Abstract

The southern sea otter (*Enhydra lutris nereis*) population at San Nicolas Island, California, has been monitored annually since the translocation of 140 sea otters to the island was completed in 1990. Monitoring efforts have varied in frequency and method across years. In 2017, in accordance with the National Defense Authorization Act for Fiscal Year 2016, the U.S. Navy and the U.S. Fish and Wildlife Service formally initiated a sea otter monitoring and research plan to determine the effects of military readiness activities on the growth or decline of the southern sea otter population at San Nicolas Island. The monitoring program, at its basic level, includes quarterly seasonal surveys of population abundance, distribution, and foraging activity. This report presents data from the program with a focus on the recent three years from winter 2023 through winter (February) 2026. From 2023 to 2026, we measured an 8.1-percent per annum decrease in population abundance (95-percent confidence interval =1.1–14.6 percent), with 106 total individuals counted as of February 2026. Historically, sea otter habitat usage at San Nicolas Island was concentrated on the west end of the island. Between 2017 and 2019, we observed increased seasonal usage of the north and south sides of the island, and in 2020-2022, a large (approximately 30-40 individuals) group of sea otters (raft) took up residence off the east end. During 2023-2026 the east end raft disappeared, and sea otters returned to their historical habitat usage patterns at the west end of the island. Foraging data were collected from summer 2023 to winter 2026 on a total of 461 foraging dives in 32 foraging bouts, and the majority of identified prey on successful dives (n=325) were sea urchins (124) followed by snails (48), bivalves (41) and crabs (23). One lobster and one octopus were also identified among the sea otter prey items. We combined these data with data from 2020-2022 to estimate overall energy intake rates that averaged 7.7 kilocalories per minute (95-percent credible interval =6.6–9.1 kilocalories per minute). These results can be useful to the planning of future monitoring and research of sea otters at San Nicolas Island.

## Introduction

The southern sea otter (*Enhydra lutris nereis*), also known as the California sea otter, is a subspecies of sea otter (*Enhydra lutris*) native to California. Before the fur trade of the 18^th^ and 19^th^ centuries, an estimated 150,000 to 300,000 sea otters ranged along the continental shelf of the northern Pacific Ocean from Japan through Mexico (Bodkin, 2015). Sea otters were hunted extensively during the fur trade, and by the time of their protection under the North Pacific Fur Seal Treaty of 1911, they were nearly extinct. The California population was believed to be extirpated until a remnant group of about 50 sea otters was discovered near Big Sur, California. The southern sea otter population is listed as threatened under the Endangered Species Act of 1973 (ESA) and considered depleted under the Marine Mammal Protection Act of 1972 (MMPA). The surviving California population of sea otters gradually increased in range and number, reaching about 1,500 individuals by the mid-1980s. However, concerns remained about the survival of this population because of its limited distribution, particularly if an oil spill or other disaster were to occur. The U.S. Fish and Wildlife Service (USFWS) developed a program to reintroduce up to 250 southern sea otters to San Nicolas Island (SNI), California, as a reserve population (USFWS, 2012). Despite the translocation of 140 sea otters between 1987 and 1990, fewer than 20 sea otters were counted at SNI for nearly a decade (table 1). Most of the translocated animals disappeared, but many returned to the mainland population and some moved to and stayed in other areas (Rathbun and others, 2000; Hatfield 2005). From the beginning of the translocation, the USFWS and later the National Biological Survey and the U.S. Geological Survey (USGS) monitored the status and trends of this population. Spring and fall surveys occurred in all years, with the spring survey at SNI coinciding within several weeks of the mainland California sea otter census and up to four seasonal surveys occurring in some of these years.

**Table 1.** Counts of independent sea otters and pups at San Nicolas Island, California, during spring surveys from 1990 through 2025. The spring 2026 survey was not completed by the time this report was prepared. * Spring surveys were cancelled during 2020 and 2021 due to the coronavirus disease 2019 pandemic restrictions.

| Year | Independents | Pups | Total |
| --- | --- | --- | --- |
| 1990 | 14 | 3 | 17 |
| 1991 | 14 | 2 | 16 |
| 1992 | 10 | 2 | 12 |
| 1993 | 7 | 4 | 11 |
| 1994 | 10 | 4 | 14 |
| 1995 | 9 | 4 | 13 |
| 1996 | 12 | 4 | 16 |
| 1997 | 16 | 0 | 16 |
| 1998 | 12 | 2 | 14 |
| 1999 | 18 | 3 | 21 |
| 2000 | 21 | 2 | 23 |
| 2001 | 21 | 5 | 26 |
| 2002 | 22 | 5 | 27 |
| 2003 | 33 | 5 | 38 |
| 2004 | 27 | 4 | 31 |
| 2005 | 22 | 3 | 25 |
| 2006 | 36 | 5 | 41 |
| 2007 | 26 | 4 | 30 |
| 2008 | 22 | 3 | 25 |
| 2009 | 27 | 6 | 33 |
| 2010 | 38 | 7 | 45 |
| 2011 | 44 | 6 | 50 |
| 2012 | 48 | 10 | 58 |
| 2013 | 54 | 8 | 62 |
| 2014 | 59 | 9 | 68 |
| 2015 | 54 | 7 | 61 |
| 2016 | 92 | 12 | 104 |
| 2017 | 72 | 9 | 81 |
| 2018 | 81 | 14 | 95 |
| 2019 | 109 | 12 | 121 |
| 2020* |  |  |  |
| 2021* |  |  |  |
| 2022 | 119 | 2 | 121 |
| 2023 | 134 | 12 | 146 |
| 2024 | 94 | 19 | 113 |
| 2025 | 111 | 14 | 125 |

The U.S. Navy (USN) owns SNI and supported the translocation of sea otters to the island with a provision that defense-related actions were exempt from ESA consultation requirements (Public Law 99-625). After the USFWS declared the translocation program a failure in 2012, additional provisions for military readiness activities were legislated through the National Defense Authorization Act (NDAA) for Fiscal Year 2016. As required by the NDAA, the USN and the USFWS, in coordination with the USGS, developed a monitoring and research plan for the study of southern sea otters in the military readiness areas of SNI and San Clemente Island, also owned by the USN, which are both among the California Channel Islands (appendix 1). Of the two islands, sea otters are currently known to occupy only SNI. Therefore, all monitoring efforts within this program were focused at SNI.

The military readiness area monitoring and research plan aims to “monitor interactions between military readiness activities and the sea otter population” (appendix 1). The plan defines three tiers of monitoring, each corresponding to an increasing management response as predefined thresholds indicating potential impacts to the SNI sea otter population from military readiness activities are exceeded. The population is currently monitored in accordance with tier 1 (“basic monitoring and research”), in which the objectives are to (1) monitor and analyze population trends at SNI, (2) monitor and analyze subtidal benthic communities and assess impacts of sea otter recovery on food dynamics, and (3) determine factors affecting population change, including relative contributions of density-dependent factors (changes in per-capita prey abundance) and density-independent factors (for example, shark mortality, military readiness activities) to variation in growth rates. The plan prescribes a strategy in which any new military readiness activities that could affect sea otters would trigger increased (tier 2) monitoring (“operational monitoring”). Additional (tier 3) monitoring (“advanced monitoring”) is triggered if, in addition to new military activities, one of the following three events occurs: (1) a dead, moribund, or stranded sea otter is found with injuries consistent with military activities is detected; (2) the population trend decreases by more than 10 percent from the average trend over the preceding 3-year period for at least 2 consecutive years for reasons unattributable to density dependence; or (3) the total sea otter population drops below 75.

The USGS addresses the objectives of tier 1 by doing quarterly sea otter population and foraging surveys and semi-annual subtidal kelp community surveys consistent with long-term monitoring programs and by developing quantitative models that describe patterns of population change. This report presents information and analyses of data collected from the quarterly seasonal sea otter surveys from July 2023 through February 2026 and is an update to earlier reports (Yee and others, 2023, Yee and others, 2020) that covered the seasonal sea otter surveys since February 2017, the time of initiation of the military readiness area monitoring plan. New information collected from continuing surveys of the long-term subtidal kelp monitoring program are presented in a separate report (Kenner and Tomoleoni, 2021). Herein, we evaluate population abundance/distribution and foraging activity around SNI, and we compare our findings to those from previous survey periods (2003–2006, 2017–2019, and 2020-2022) when comparable types of monitoring were done. We present sea otter density maps with annual imagery snapshots of kelp distribution around SNI. We also analyze foraging success, including prey compositions and energy intake.

## Methods

In this section, we describe the protocols we used at SNI to survey and analyze sea otter population abundance and spatial distribution and sea otter foraging activity and outcomes.

### Population Abundance and Distribution Surveys

For this reporting period, 2023–2026, winter surveys occurred in January/February, spring surveys in April/May, summer surveys in July/August, and fall surveys in December (as weather conditions and island access permitted). Surveys were completed following protocols similar to the annual range-wide southern sea otter census in California (Hatfield and others, 2019), except there were usually two or three attempts to obtain a daily count of the entire island on different days of the survey trip whereas an extensive count of mainland California is obtained once annually over multiple days as synchronously as possible. For each survey day, observers were assigned to a section of the island comprising one or more of nine survey areas (fig. 1). Each observer was equipped with a high resolution 50-80X spotting scope (Questar Inc., New Hope, Pennsylvania) and binoculars (10X; various brands). The observers started at one end of their assigned section and selected an observation point that provided a good vantage point of nearby sea otter habitat. From the vantage point, the observers scanned with unaided eyes and with binoculars for sea otters or objects that were suspected to be sea otters. Large groups, suspicious objects, and distant habitat were scanned with the spotting scope. After sufficient time (depending on viewing conditions, extent of area surveyed, and sea otter abundance) elapsed to make a thorough count of all sea otters within this first segment of habitat, the observer moved to another location that provided good viewing of the next segment of habitat, contiguous with the first segment. This process was repeated until the entire section was counted; in most cases observers used the same counting locations year after year for consistency.

**Figure 1.**
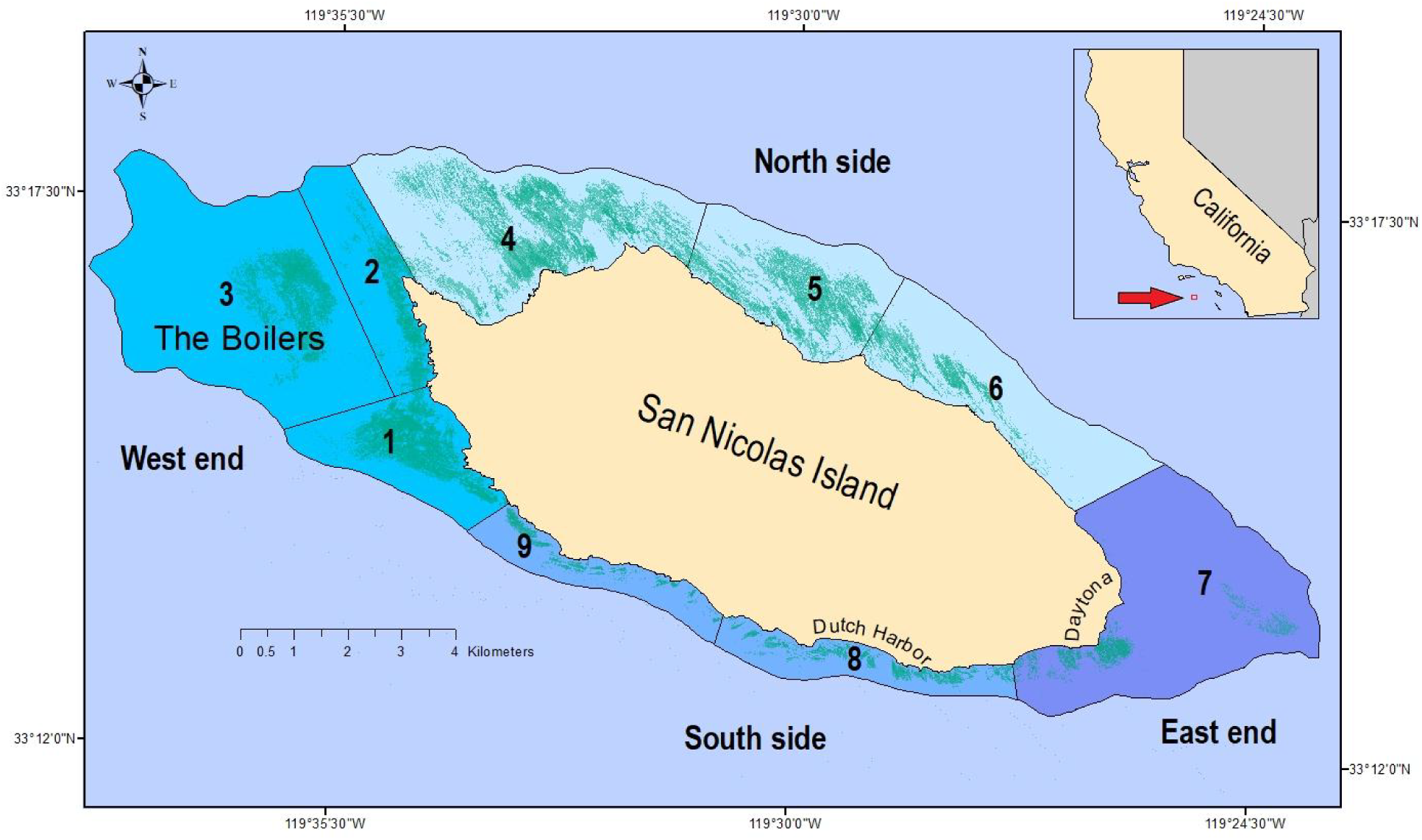
Sea otter habitat within a 30-meter isobath around San Nicolas Island, California. Sea otter counts are tallied into nine contiguous survey areas numbered from 1 to 9 and grouped into 4 sections (blue shades). Mapped kelp canopies (shaded dim green) are derived and merged from aerial imagery taken in the fall seasons of 2017–2019.

All sea otter observations were marked as points directly onto paper maps that showed major stationary features of the area (shoreline configuration and offshore rocks) for reference. Kelp beds from an earlier year were displayed on these map sheets and could be used as a secondary reference due to their tendency to persist; however, surface kelp canopy contracts, expands, and shifts amongst months and years, and these printed images were not necessarily reliable for marking the precise locations of sea otters for comparisons across surveys. These features, regardless, allowed sea otter locations to be mapped relative to available visual cues within the same survey, which reduced the chances of double-counting or undercounting when the observer moved to the next observation point.

To minimize double-counting or detection biases caused by sea otter movements, each total island count included surveys of all nine areas of the island on one survey day, and attempts were made to avoid double-counting active sea otters that were observed swimming towards the next adjacent segment. In addition to the sea otter locations, associated data recorded directly onto the maps included the number of independent (non-pup) sea otters, number and age class of pups (pups are classed as either small or large depending on the presence or absence of the natal pelage, relative size, and behavior), behavior (resting, foraging, grooming, swimming, mating, or other), group size, and micro-habitat type (open water or kelp). Time and general survey conditions also were recorded. Viewing conditions were rated from excellent to poor (coded 4 to 0). Shore-based surveys are estimated to detect about 90–95 percent of the sea otters located in the nearshore area (Estes and Jameson, 1988), but distances exceeding 1 kilometer (km) offshore and factors such as viewing conditions and sea otter grouping and behaviors could influence detectability. Surveys were repeated on multiple days during each survey trip, and the maximum one-day total was taken as the island-wide count to minimize detection biases.

### Ecosystem Monitoring

Subtidal kelp forest monitoring has occurred semi-annually in April and October in most years since October 1980, and information on these surveys through fall 2019 is reported in detail in a separate report (Kenner and Tomoleoni, 2021). Subtidal monitoring was suspended in 2020 and 2021 due to enforced distancing measures on vessels resulting from the coronavirus disease of 2019 (COVID-19) pandemic but was resumed in 2022. In addition to subtidal kelp forest monitoring, we obtained multi-spectral imagery of the survey areas around SNI contracted by the U.S. Navy and taken by staff at Ocean Imaging Corporation (http://www.oceani.com, Littleton, Colorado, formerly also known as Atoll Ventures) annually during the fall of 2014 through 2019. These images were taken at 0.38-meter resolution in red, green, blue, and near-infrared bands, and pixels were classified into exposed or submerged giant kelp canopies based on supervised learning algorithms. We obtained raster and shapefiles of these images and used geographic information systems (GIS) to develop maps of sea otter distributions relative to areas with kelp.

### Foraging Surveys

Foraging data were collected opportunistically during the same seasonal surveys when population abundance and distribution data were collected. Observation methods followed well-established protocols (Estes and others, 1981; Ralls and others, 1995; Tinker and others, 2008). Field observations were collected during daylight hours by teams of 1–2 observers searching for actively foraging sea otters near enough to shore where they could be visually distinguished from other sea otter individuals and prey was possible to identify using a 50-80X spotting scope. Foraging bouts, defined as continuous sequences of feeding dives made by the focal sea otter, were observed until data from at least 20 dives in the bout were recorded, visual contact with the focal sea otter was lost, or the sea otter stopped foraging. The information recorded during these bouts included date and time, location of the bout, duration of the subsurface dive interval (dive time) and the post-dive surface interval (surface time) for each feeding dive (in seconds), outcome of each dive (in other words, whether or not prey was captured), type of prey captured, number and size of prey items, per-item handling time (number of seconds required to handle and consume each item), whether or not tools were used to handle the prey, and ambient conditions (including sea-state, wind, and so forth). Prey size was recorded as the estimated diameter of the shell or maximum body dimension (excluding appendages), categorized into 5- centimeter incremented size-classes. Each 5-centimeter increment represents one typical adult sea otter paw-width and was subdivided further into thirds within that size-class for increased precision. For observations where prey could not be reliably identified to species, the items in question were assigned to the lowest possible taxonomic level. Any items that could not be reliably categorized to any taxonomic level were classified as unknown prey, with size class and number of items still recorded if possible. Additional information recorded by observers included numbers of prey items that were carried over from a previous dive or stolen by or from the focal animal, proportion consumed in cases where prey was only partially eaten, and, in the case of females with dependent pups, the amounts of these prey that were shared with the pups.

### Analyses

#### Abundance and Trends

Using all seasonal counts, we graphically modeled trends in island-wide sea otter abundance using nonlinear regressions fitted by locally estimated sum of squares (LOESS). LOESS trends were fitted locally by the “geom_smooth” function of the “ggplot2” package in R statistical software using a default span of 0.75 of data observations (Wickham 2016; R Core Team, 2025). We used generalized linear models (GLM) on 3-year moving intervals to estimate variable rates of population growth across time, beginning with the 1990–1993 interval and ending with the 2023–2026 interval. We modeled sea otter counts (*C_t_*, for year *t*) within each interval by fitting a linear trend where replicate counts *C_t_* vary according to an overdispersed Poisson model with mean *μ_t_* = *exp*(*β_0_* + *βt*) and variance *cμ_t_*. The multiplier (*c*) in the variance represents an overdispersion factor, or in other words, the amount by which the variance of count data exceeds that of a standard Poisson distribution, in which both the mean and the variance are *μ_t_* (McCullagh and Nelder, 1989). We used the estimated slope coefficient (*β*) to calculate and test multiplicative rates of annual change in population size, *λ* = *exp*(*β*), based on the mathematical equivalence between *exp*(*β*) and the ratio in expected counts between two consecutive years, that is *μ_t_*_+1_/*μ_t_*. For the 2017–2026 years, we included replicate counts from all available seasons (*C_t_*_,*s*_, for season *s*) and tested for seasonal differences in island-wide total counts. We calculated the pup-to-independent ratio by summing the counts of sea otter pups across surveys and dividing by the corresponding sum of counts of independent sea otters, and we multiplied this ratio by 100 to derive an index of the number of sea otter pups per 100 independent sea otters. We used R statistical software to complete all statistical analyses and the “ggplot2” package to create graphics (Wickham 2016; R Core Team, 2025).

#### Distributions

We used ArcGIS Pro software (Environmental Systems Research Institute, Inc., Redlands, California, https://www.esri.com/, version 3.5) to illustrate and interpret the spatial distribution of sea otter locations around San Nicolas Island for the years 2023–2026. We modeled spatial distributions by aggregating sea otter locations from the high-count dates across multiple survey trips, and we used kernel density estimation techniques with a 2-km smoothing parameter to map spatially explicit estimates of relative sea otter density. We evaluated this density map for the aggregation of 11 seasonal surveys from spring (April) 2023 through winter (February) 2026, and we compared that to the previous density maps for the aggregation of 12 seasonal surveys completed from February 2017 through November 2019 and for the aggregation of 8 seasonal surveys completed from February 2020 through October 2022. We used descriptive bar charts that represent the periphery around SNI to visually assess changes in sea otter distributions among the nine areas. For statistically analyzing changes in distribution, we grouped the nine areas into four sections of the island (west end, north side, east end, and south side; Figure 1) to ensure adequate sample sizes within each area category, and we used GLM to test for significant changes in distributions among periods (2017–2019, 2020–2022, and 2023–February 2026) and among years and seasons within the recent period from 2023 to February 2026. We constructed additional density maps to examine spatial differences when any of these factors were found to be significant.

#### Foraging

We combined the foraging data from 2023–February 2026 and categorized them into seasons (winter, spring, summer, and fall), and sex (males, females, and unknown), and we report foraging outcomes by these categories. We also examined prey composition within these categories in terms of the proportion of items within eight prey classes: (1) abalone, (2) bivalves, (3) crabs, (4) lobsters, (5) octopus, (6) snails, (7) urchins, and (8) other. For the assessments of prey composition and energy intake rates, the sample sizes of the foraging data collected during 2023–February 2026 were too small to separate estimates by year. For relative frequencies of observed and identified prey classes, we combined foraging data from the years 2023–February 2026 and compared them to data from individual years from 2003 to 2006 and 2017 to 2019, when sample sizes per year were robust, and from 2020–2022 combined, when sample sizes per year were limited. For the analyses of prey compositions based on biomass and for energy intake rates, we combined foraging data from the years 2020–February 2026 and compared with those of the grouped years 2003–2006 and of the separate years from 2017 to 2019. To obtain quantitative measures of the composition of various prey species in each individual’s diet and energy intake rates (kilocalories per minute), we converted field counts of prey capture frequency and prey shell diameter to estimates of consumed biomass and caloric content using species-specific power functions for converting prey diameter to wet edible biomass and kilocalories per gram (Oftedal and others, 2007). These data were then analyzed using a Monte-Carlo simulation procedure to estimate diet composition (in terms of the proportion of consumed biomass contributed by each prey group) and rate of energy gain, or kilocalories consumed per minute of feeding (Tinker and others, 2008). The Monte-Carlo procedure incorporates sampling uncertainty and adjusts for recognized biases associated with direct observations of sea otter foraging; a detailed description of the algorithm is provided elsewhere (Tinker and others, 2012; Tinker, 2015). We also used the Shannon-Wiener index to compare diet diversity based on simulated estimates of prey type biomass between the 2020–February 2026 period and those of the previous periods, 2003–2006 and 2017–2019 (Tinker and others, 2008).

## Results

### Abundance and Trends

Shortly after the translocation of 140 sea otters to SNI from 1987–1989, most disappeared from SNI, leaving a population of fewer than 20 sea otters during the majority of the first decade (table 1). However, this population gradually grew to exceed 100 individuals by the time of the spring 2016 survey and continued to exceed 100 individuals in every survey since fall 2018 except winter 2024 (96 individuals recorded) and fall 2025 (95 individuals recorded) (table 2, fig. 2). Counts from 2024 through February 2026 ranged between 95 and 125 total sea otters (table 2; fig. 3).

**Figure 2.**
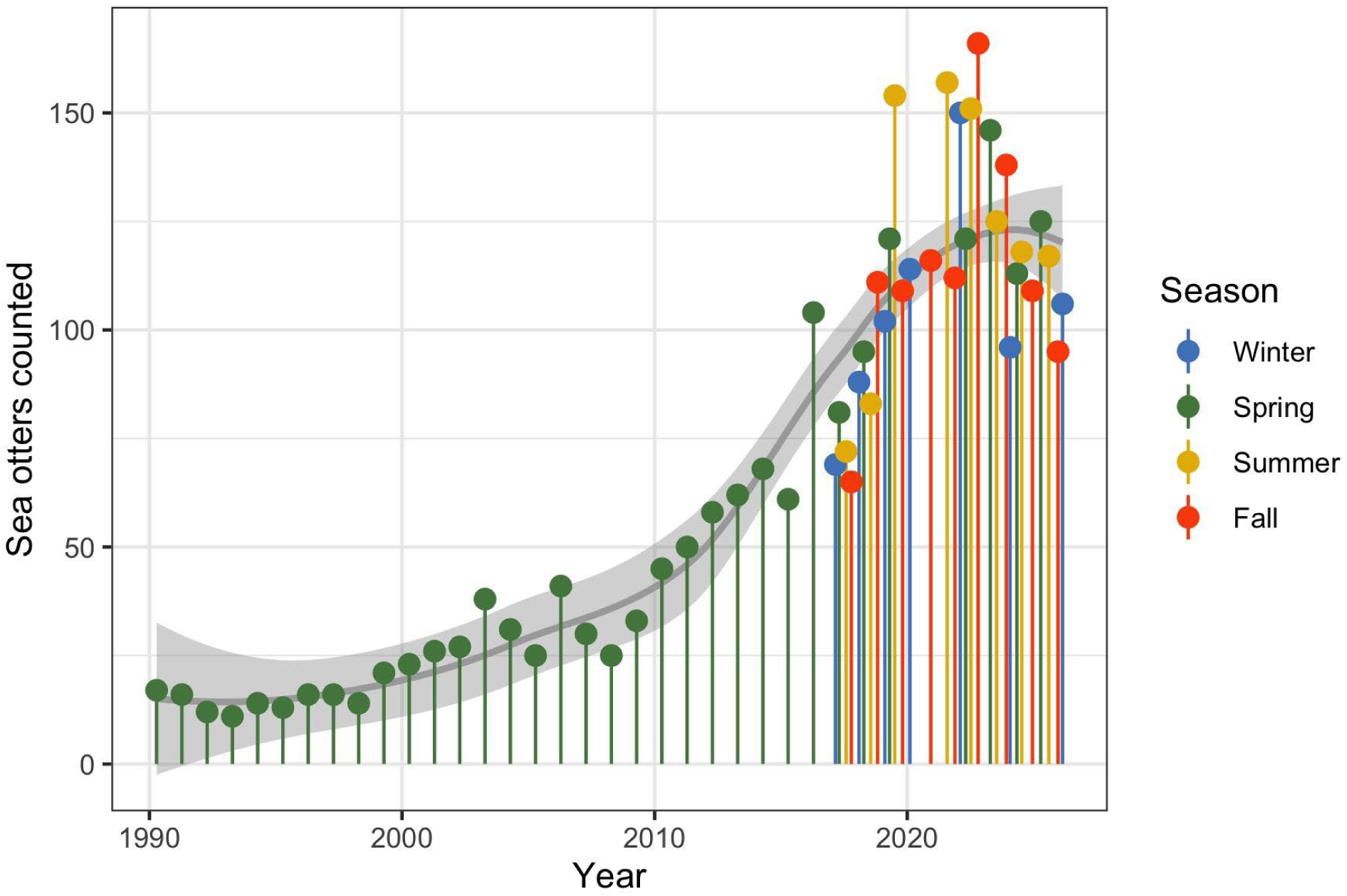
Number of sea otters (independent + pups) counted at San Nicolas Island, California, from 1990 to 2026. Seasonal monitoring for the Navy began in 2017. Solid line represents a smooth regression of the total sea otter counts in relationship to year, fitted by the locally estimated sum of squares (LOESS) method, and shaded band represents 95 percent confidence intervals around the fitted estimates.

**Figure 3.**
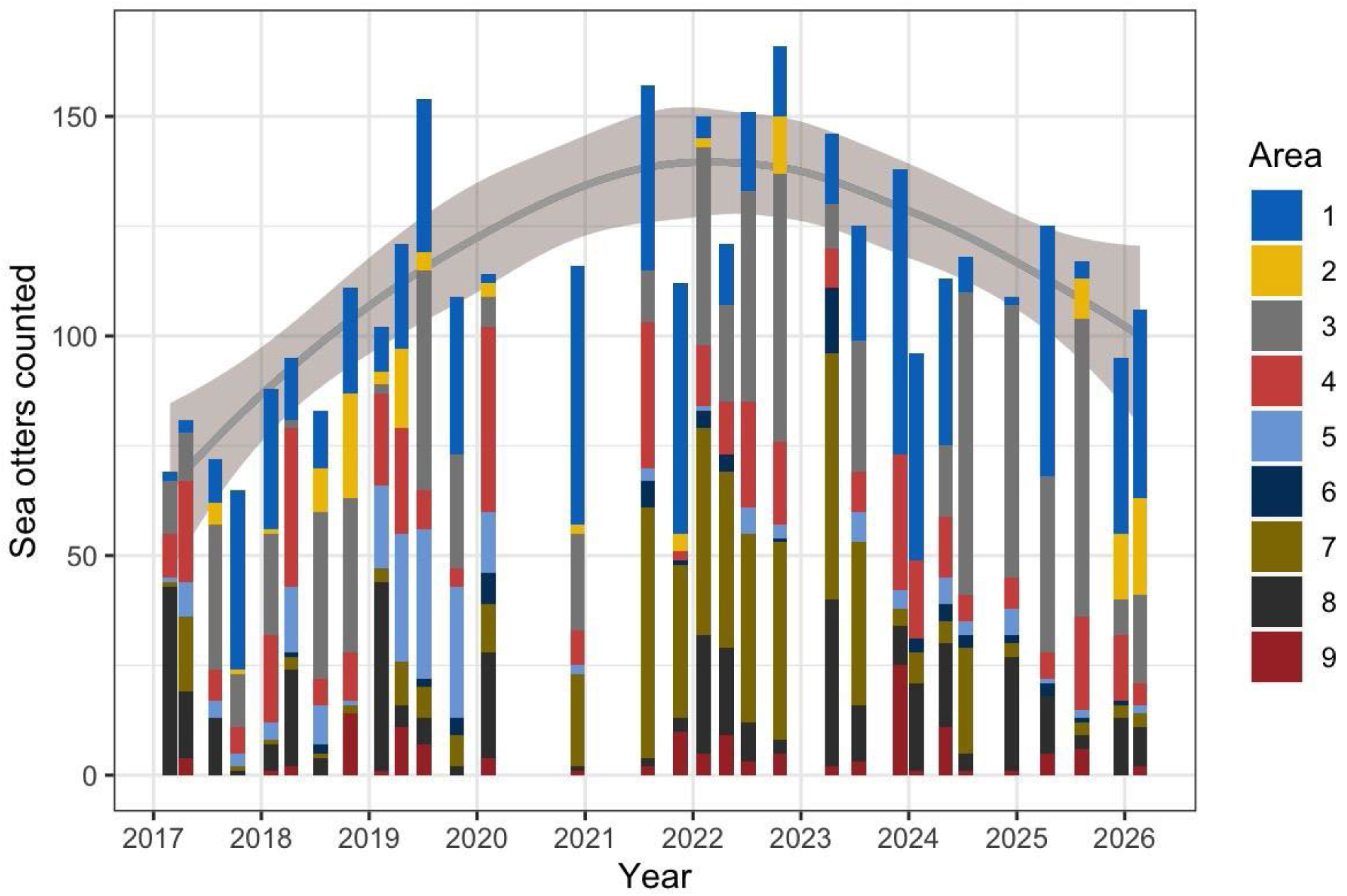
Number of sea otters counted at each of the 9 areas at San Nicolas Island, California, during 31 seasonal surveys from February 2017 to February 2026. Solid gray curve represents a smooth regression of the total sea otter counts in relationship to year, fitted by the locally estimated sum of squares (LOESS) method (gray line), and light gray-shaded with 95-percent confidence intervals. The LOESS regression was smoothed using a 0.75 proportion of the plotted data localized around each year. Count data are presented in table 2.

**Table 2.** Count of independent sea otters and pups at San Nicolas Island, California, during seasonal surveys from winter 2017 through winter 2026, by area and combined for the entire island.

| Season | Year | Month | Area count (independents + pups) |  |  |  |  |  |  |  |  | Total Island count |  |  |
| --- | --- | --- | --- | --- | --- | --- | --- | --- | --- | --- | --- | --- | --- | --- |
|  |  |  | 1 | 2 | 3 | 4 | 5 | 6 | 7 | 8 | 9 | Independents | Pups | Total |
| Winter | 2017 | 2 | 2 + 0 | 0 + 0 | 12 + 0 | 9 + 1 | 1 + 0 | 0 + 0 | 1 + 0 | 37 + 6 | 0 + 0 | 62 | 7 | 69 |
| Spring | 2017 | 4 | 3 + 0 | 0 + 0 | 11 + 0 | 20 + 3 | 8 + 0 | 0 + 0 | 13 + 4 | 13 + 2 | 4 + 0 | 72 | 9 | 81 |
| Summer | 2017 | 7 | 10 + 0 | 5 + 0 | 31 + 2 | 5 + 2 | 4 + 0 | 0 + 0 | 0 + 0 | 9 + 4 | 0 + 0 | 64 | 8 | 72 |
| Fall | 2017 | 10 | 36 + 5 | 1 + 0 | 9 + 3 | 6 + 0 | 3 + 0 | 0 + 0 | 1 + 0 | 1 + 0 | 0 + 0 | 57 | 8 | 65 |
| Winter | 2018 | 2 | 30 + 2 | 1 + 0 | 18 + 5 | 19 + 1 | 4 + 0 | 0 + 0 | 1 + 0 | 6 + 0 | 1 + 0 | 80 | 8 | 88 |
| Spring | 2018 | 4 | 12 + 2 | 0 + 0 | 2 + 0 | 31 + 5 | 15 + 0 | 1 + 0 | 2 + 1 | 16 + 6 | 2 + 0 | 81 | 14 | 95 |
| Summer | 2018 | 7 | 12 + 1 | 10 + 0 | 33 + 5 | 5 + 1 | 9 + 0 | 2 + 0 | 1 + 0 | 4 + 0 | 0 + 0 | 76 | 7 | 83 |
| Fall | 2018 | 10 | 22 + 2 | 24 + 0 | 32 + 3 | 10 + 1 | 1 + 0 | 0 + 0 | 2 + 0 | 0 + 0 | 9 + 5 | 100 | 11 | 111 |
| Winter | 2019 | 2 | 8 + 2 | 2 + 1 | 2 + 0 | 18 + 3 | 19 + 0 | 0 + 0 | 3 + 0 | 32 + 11 | 1 + 0 | 85 | 17 | 102 |
| Spring | 2019 | 4 | 20 + 4 | 17 + 1 | 0 + 0 | 23 + 1 | 29 + 0 | 0 + 0 | 8 + 2 | 4 + 1 | 8 + 3 | 109 | 12 | 121 |
| Summer | 2019 | 7 | 32 + 3 | 4 + 0 | 45 + 5 | 9 + 0 | 31 + 3 | 2 + 0 | 6 + 1 | 6 + 0 | 6 + 1 | 141 | 13 | 154 |
| Fall | 2019 | 10 | 29 + 7 | 0 + 0 | 26 + 0 | 4 + 0 | 29 + 1 | 4 + 0 | 6 + 1 | 2 + 0 | 0 + 0 | 100 | 9 | 109 |
| Winter | 2020 | 2 | 2 + 0 | 3 + 0 | 6 + 1 | 37 + 5 | 13 + 1 | 7 + 0 | 11 + 0 | 20 + 4 | 3 + 1 | 102 | 12 | 114 |
| Fall | 2020 | 12 | 47 + 12 | 2 + 0 | 19 + 3 | 7 + 1 | 1 + 1 | 0 + 0 | 21 + 0 | 1 + 0 | 1 + 0 | 99 | 17 | 116 |
| Summer | 2021 | 7 | 37 + 5 | 0 + 0 | 12 + 0 | 31 + 2 | 3 + 0 | 6 + 0 | 57 + 0 | 2 + 0 | 2 + 0 | 150 | 7 | 157 |
| Fall | 2021 | 11 | 47 + 10 | 4 + 0 | 0 + 0 | 2 + 0 | 0 + 0 | 1 + 0 | 35 + 0 | 2 + 1 | 7 + 3 | 98 | 14 | 112 |
| Winter | 2022 | 2 | 5 + 0 | 2 + 0 | 39 + 6 | 13 + 1 | 1 + 0 | 4 + 0 | 47 + 0 | 21 + 6 | 5 + 0 | 137 | 13 | 150 |
| Spring | 2022 | 4 | 13 + 1 | 0 + 0 | 22 + 0 | 11 + 1 | 0 + 0 | 4 + 0 | 40 + 0 | 20 + 0 | 9 + 0 | 119 | 2 | 121 |
| Summer | 2022 | 7 | 17 + 1 | 0 + 0 | 43 + 5 | 21 + 3 | 5 + 1 | 0 + 0 | 43 + 0 | 7 + 2 | 3 + 0 | 139 | 12 | 151 |
| Fall | 2022 | 10 | 14 + 2 | 11 + 2 | 56 + 5 | 18 + 1 | 2 + 1 | 1 + 0 | 45 + 0 | 3 + 0 | 5 + 0 | 155 | 11 | 166 |
| Spring | 2023 | 4 | 14 + 2 | 0 + 0 | 9 + 1 | 8 + 1 | 0 + 0 | 11 + 4 | 55 + 1 | 35 + 3 | 2 + 0 | 134 | 12 | 146 |
| Summer | 2023 | 7 | 22 + 4 | 0 + 0 | 29 + 1 | 9 + 0 | 7 + 0 | 0 + 0 | 35 + 2 | 13 + 0 | 3 + 0 | 118 | 7 | 125 |
| Fall | 2023 | 12 | 57 + 8 | 0 + 0 | 0 + 0 | 30 + 1 | 4 + 0 | 0 + 0 | 4 + 0 | 8 + 1 | 20 + 5 | 123 | 15 | 138 |
| Winter | 2024 | 1 | 36 + 11 | 0 + 0 | 0 + 0 | 16 + 2 | 0 + 0 | 3 + 0 | 7 + 0 | 18 + 2 | 1 + 0 | 81 | 15 | 96 |
| Spring | 2024 | 5 | 30 + 8 | 0 + 0 | 15 + 1 | 11 + 3 | 6 + 0 | 3 + 1 | 4 + 1 | 16 + 3 | 9 + 2 | 94 | 19 | 113 |
| Summer | 2024 | 7 | 7 + 1 | 0 + 0 | 65 + 4 | 6 + 0 | 3 + 0 | 3 + 0 | 24 + 0 | 4 + 0 | 1 + 0 | 113 | 5 | 118 |
| Fall | 2024 | 12 | 2 + 0 | 0 + 0 | 62 + 0 | 6 + 1 | 6 + 0 | 2 + 0 | 3 + 0 | 19 + 7 | 1 + 0 | 101 | 8 | 109 |
| Spring | 2025 | 4 | 49 + 8 | 0 + 0 | 37 + 3 | 5 + 1 | 1 + 0 | 2 + 1 | 0 + 0 | 13 + 0 | 4 + 1 | 111 | 14 | 125 |
| Summer | 2025 | 8 | 4 + 0 | 9 + 0 | 64 + 4 | 19 + 2 | 2 + 0 | 1 + 0 | 3 + 0 | 3 + 0 | 6 + 0 | 111 | 6 | 117 |
| Fall | 2025 | 12 | 32 + 8 | 12 + 3 | 8 + 0 | 15 + 0 | 0 + 0 | 1 + 0 | 3 + 0 | 12 + 1 | 0 + 0 | 83 | 12 | 95 |
| Winter | 2026 | 2 | 37 + 6 | 19 + 3 | 20 + 0 | 5 + 0 | 2 + 0 | 0 + 0 | 3 + 0 | 8 + 1 | 2 + 0 | 96 | 10 | 106 |

The sea otter count, as of the most recent winter survey (February 2026), was 106, which included 96 independents and 10 pups, and the estimated rate of population growth based on all seasonal surveys during the 2023–2026 period was *λ*_2026_ = 0.919 (95-percent CI = 0.854– 0.989), corresponding to a 8.1-percent annual decline during this period (95-percent CI = 1.1– 14.6 percent). Annual population growth rates varied during the past three decades, with about half of the estimates and 95-percent CI exceeding 1, indicating statistically positive annual growth, and most growth occurring after 2010 (figs. 2, 4). However, the trends from the three most recent 3-year moving intervals (2021–2024, 2022–2025, and 2023–2026) have each shown significant population declines (fig. 4).

**Figure 4.**
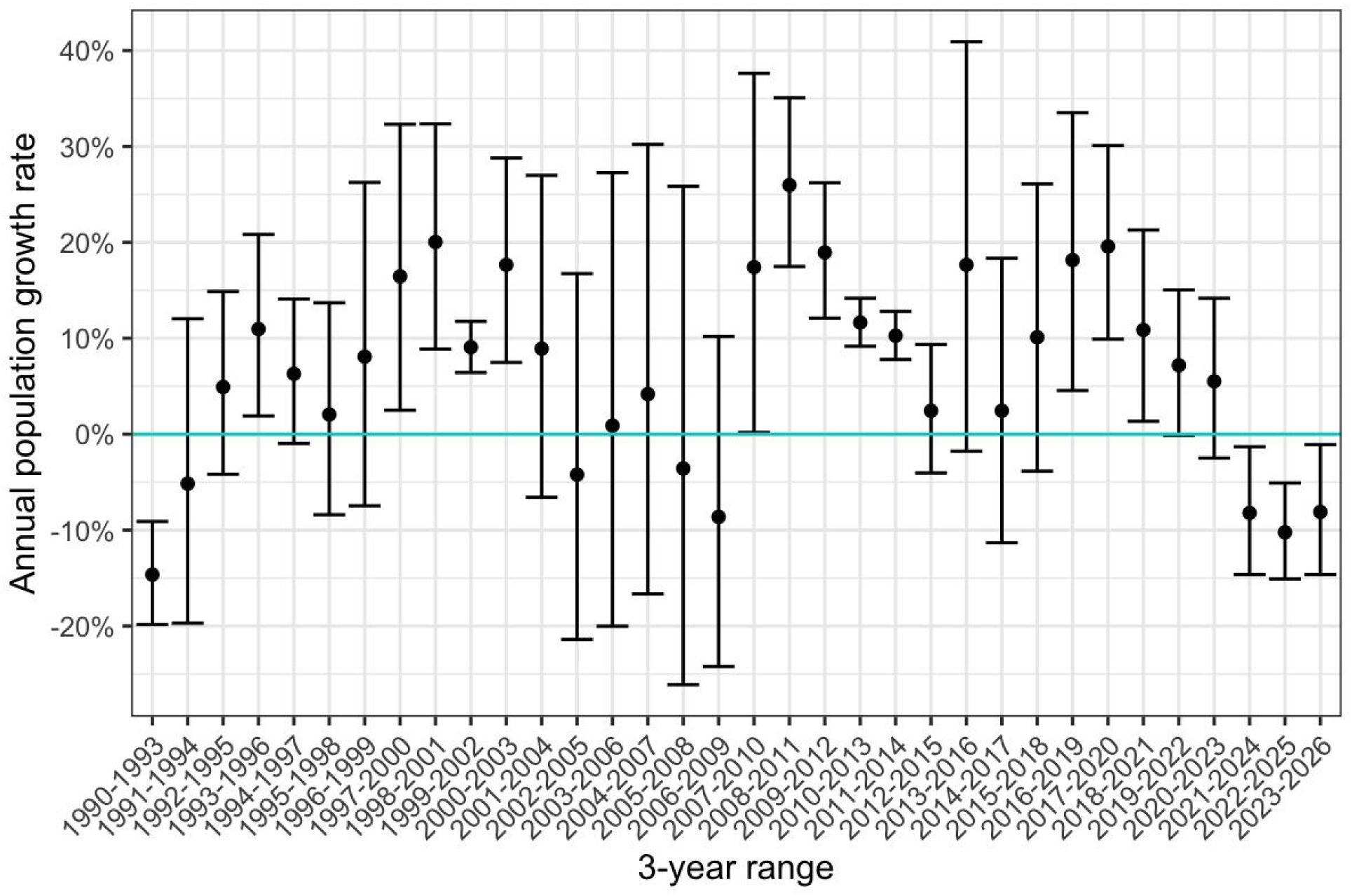
Estimated rates of annual population growth of sea otters at San Nicolas Island, California, based on seasonal surveys during 3-year moving intervals from 1990 to 2026, with 95-percent confidence intervals. Positive rates represent population growth, whereas negative rates represent population decline. Confidence intervals that overlap with 0 indicate insufficient evidence to conclude growth or decline.

The pup-to-independent ratio for surveys in the 2023–February 2026 period was 11.0 pups to 100 independent sea otters, which was higher than the pup-to-independent ratio for surveys in the previous (2020–2022) period (9.5 pups per 100 independent sea otters). Lower pup ratios are expected following periods of population growth, as more juveniles and subadults recruit into the pre-breeding population.

### Density Maps and Spatial Distributions

We compared the densities and distributions of sea otters during all surveys from the spring 2023–winter 2026 period with those of the prior time periods (2017–2019, 2020–spring 2023) previously reported by Yee and others (2023). Sea otter locations were aggregated across all surveys during each period to develop kernel density surfaces representing the relative density of cumulative counts (fig. 5). In these maps, sea otters appear to have shifted in distribution around the island through time. Prior to 2017 (not pictured) sea otters primarily used the west end of SNI, with some number of sea otters found on the south side but few or no sea otters found on the north side and east end. From 2017 to 2019, sea otter use of the island was the most widespread, with considerable usage of the north side of the island. Between 2020 and 2022, a large group of more than 40 sea otters became established off the east end of SNI, and the distribution was mostly split between the west and east ends, while very few sea otters used the areas in between. From 2023 to 2026, sea otters appeared to have reestablished usage of more of the island, and higher densities have returned to the west side, particularly The Boilers (Survey Area 3) and Survey Area 1. A finer scale annual look at the 2023–2026 period revealed marked change during the past three years. Specifically, the large group of 40+ otters at the east end suddenly disappeared after the summer 2023 survey causing the sea otter usage of SNI to shift from broad island-wide usage (though with persistent patches of low concentrations on the north and south sides) to heavier usage of The Boilers and Survey Area 1 at the west end in 2024 (fig. 6A, 6B). During the summer 2024 survey, for the first time at SNI, a large group of sea otters was found in a distant offshore seasonal kelp bed at the east end (Area 7), just north of the sand spit (fig. 6B). By 2025, the westward shift became more pronounced, with high densities of sea otters occurring on the west end and extremely low densities, mainly singles and pairs of sea otters, occurring sporadically around the rest of the island (fig. 6C). Since 2025, the distribution of sea otters has more closely resembled historical distributions at SNI than any other survey period since surveys began under the Southern Sea Otter Military Readiness and Research Monitoring Plan in 2017.

**Figure 5.**
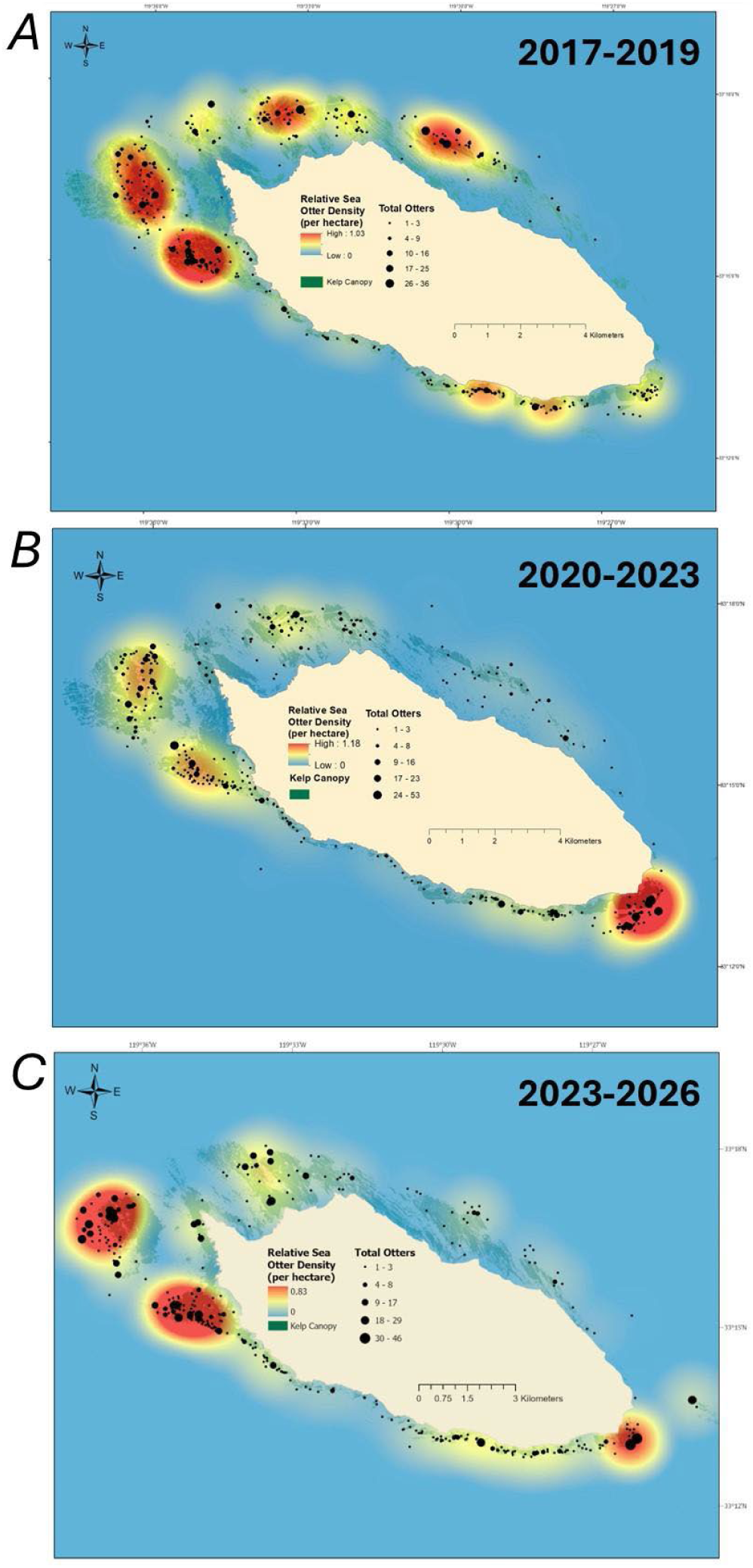
Relative sea otter density during seasonal surveys combined from *A*, winter 2017 through fall 2019 (12 surveys); *B*, winter 2020 through spring 2023 (9 surveys); and C, spring 2023 through winter 2026 (11 surveys). Sea otter locations (black dots) were aggregated across the surveys to develop kernel density surfaces representing the relative density of total counts distributed around San Nicolas Island, California. Kelp canopy images taken during the fall of 2016 through 2019 are merged and displayed for reference.

**Figure 6.**
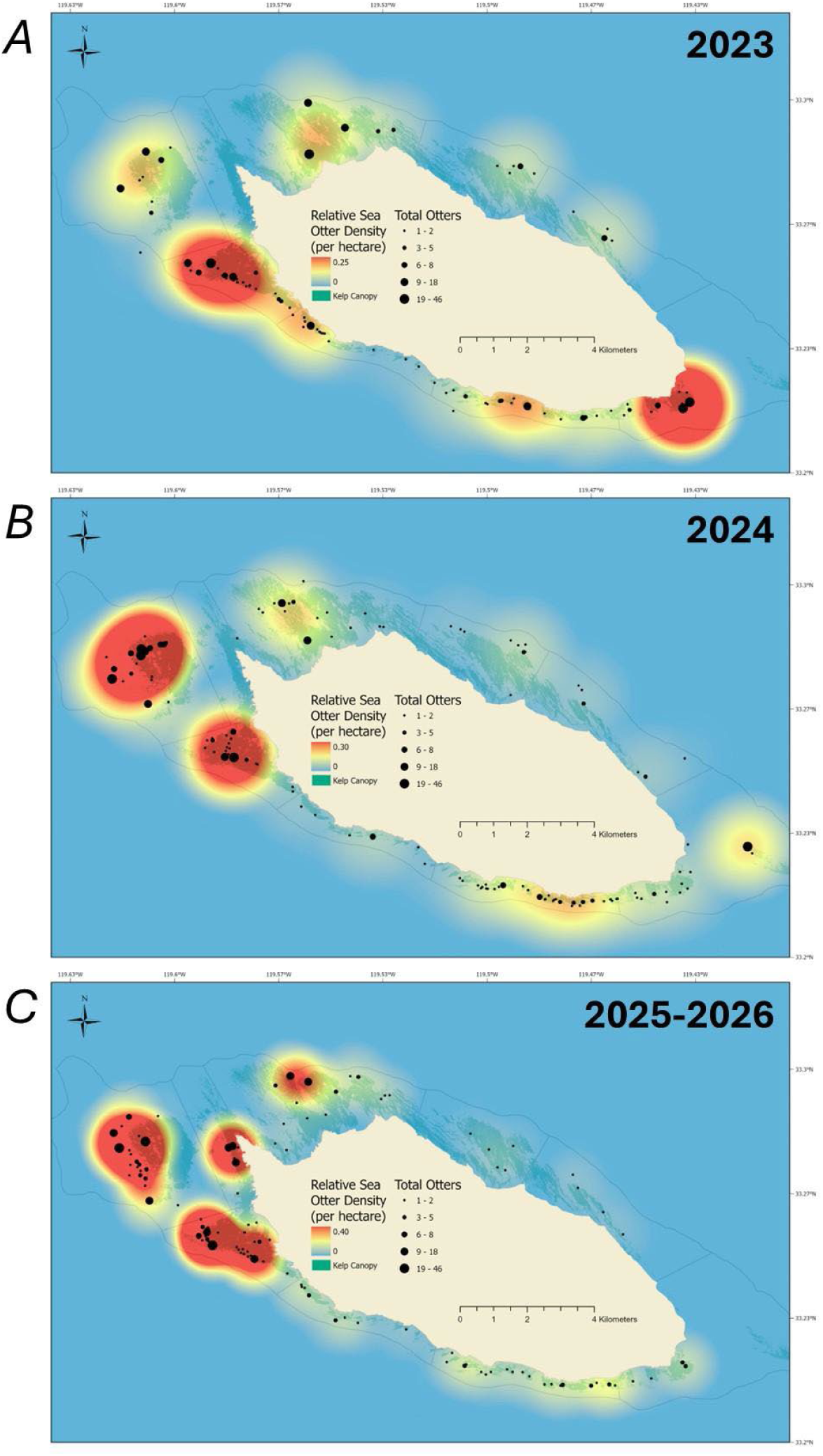
Relative sea otter density during seasonal surveys combined from *A*, 2023 (3 surveys); *B*, 2024 (4 surveys); and *C*, 2025-2026 (4 surveys). Sea otter locations (black dots) were aggregated across the surveys to develop kernel density surfaces representing the relative density of total counts distributed around San Nicolas Island, California. Kelp canopy images taken during the fall of 2017 through 2019 are merged and displayed for reference.

These maps also display sea otter densities overlaid with kelp canopy distributions derived from aerial imagery collected during the fall of 2017 to 2019. Surface kelp extent and distribution change significantly by season, with the maximum extent typically occurring in early fall. Kelp surveys were done once annually and were scheduled in the fall with the intent of capturing images of the maximum extent of kelp canopy in each year. After winter storms, the surface kelp extent is largely reduced, and the distribution changes seasonally. Thus, fall kelp distributions are not strictly representative of where kelp beds were located during seasons when sea otter surveys were completed, but they are an indicator of where kelp beds were likely to be present.

Circular plots provide a graphical summary of the distribution of sea otters counted around the perimeter of the island. The nine areas defined in figure 1 are arranged on a circular plot in the general cardinal direction as they are oriented around the island (fig.7). Most of the population distribution tends to concentrate on the west end, specifically at areas 1 and 3 (The Boilers), and the rise and fall of the east end group in Area 7 is also apparent (fig. 7).

**Figure 7.**
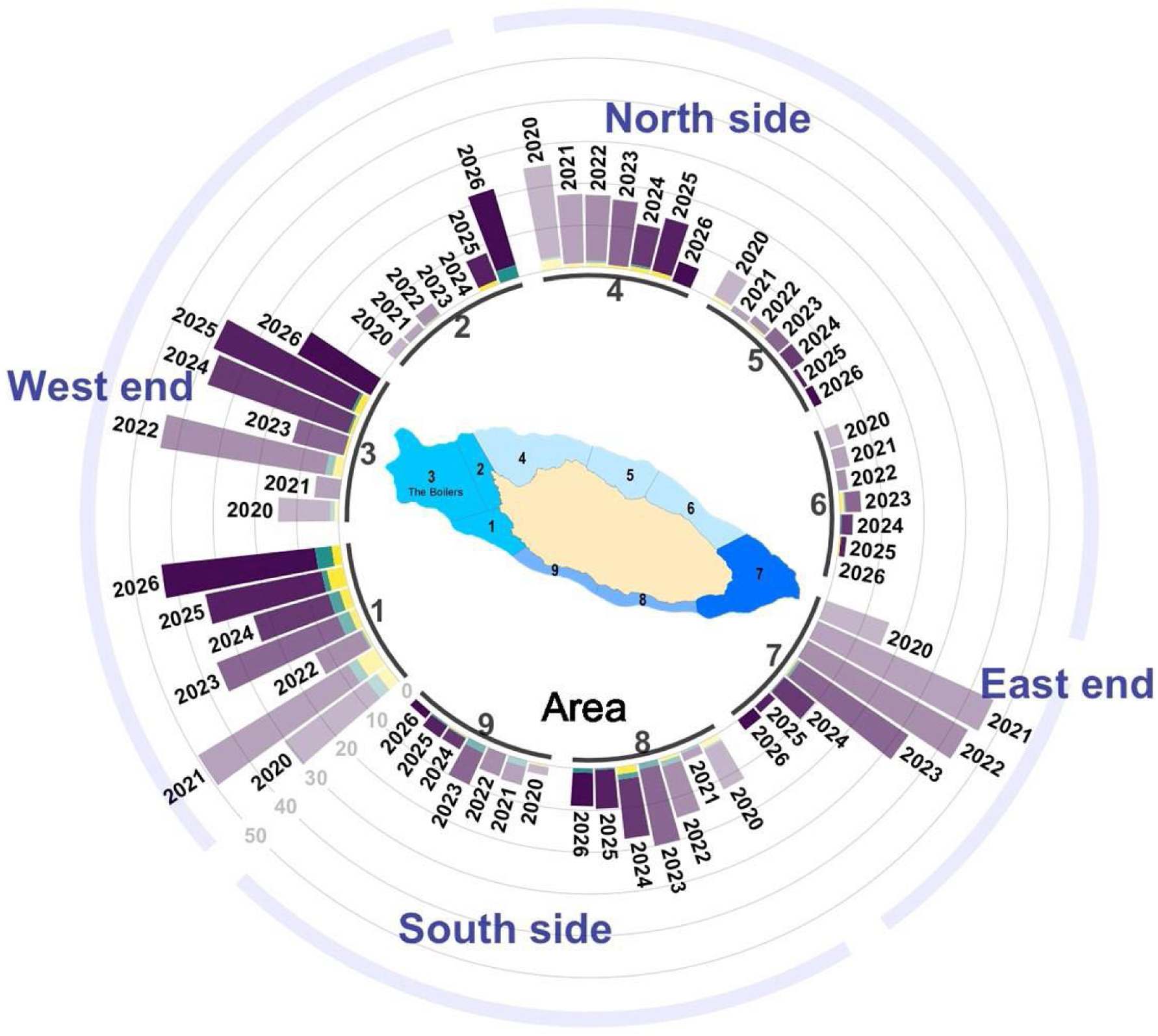
Average number of sea otters counted per year (2020-2026) at nine survey areas (refer to map inset for locations of numbered areas) around San Nicolas Island, California. Bars are grouped by area, color coded by age-size class (purple=adults, green=large pups, and yellow=small pups), and shades darken with later years (2020 is lightest, and 2026 is darkest). Rotating clockwise around the cardinal directions of the island, starting at the southwest, areas 1, 2, and 3 (The Boilers) represent the west end of the island; areas 4, 5, and 6 represent the north side; area 7 represents the east end; and areas 8, and 9 represent the south side.

Sea otter distribution around the island varied seasonally and by year during 2023-winter 2026. Areas 1 and 3 (The Boilers) on the west end consistently have the most sea otters during summer and fall, with relatively low winter densities at The Boilers, specifically, likely due to a lack of kelp canopy during the winter season (fig. 8). Spring and summer were characterized by high usage of the east end, though this seasonal distribution was heavily influenced by the presence of the large, east-end raft in spring and summer of 2023, that subsequently disappeared by the fall 2023 survey (fig. 8). The south side of the island had lower densities of sea otters overall, but counts reached a peak in winter and spring, when kelp canopy was less available on the more exposed west end of the island. Area 6 on the eastern end of the north side, where there is little to no kelp during all seasons, continued to be the least utilized region of SNI. Spatial density maps for each of the four seasons reflect these seasonal spatial distribution patterns in greater detail (fig. 8).

**Figure 8.**
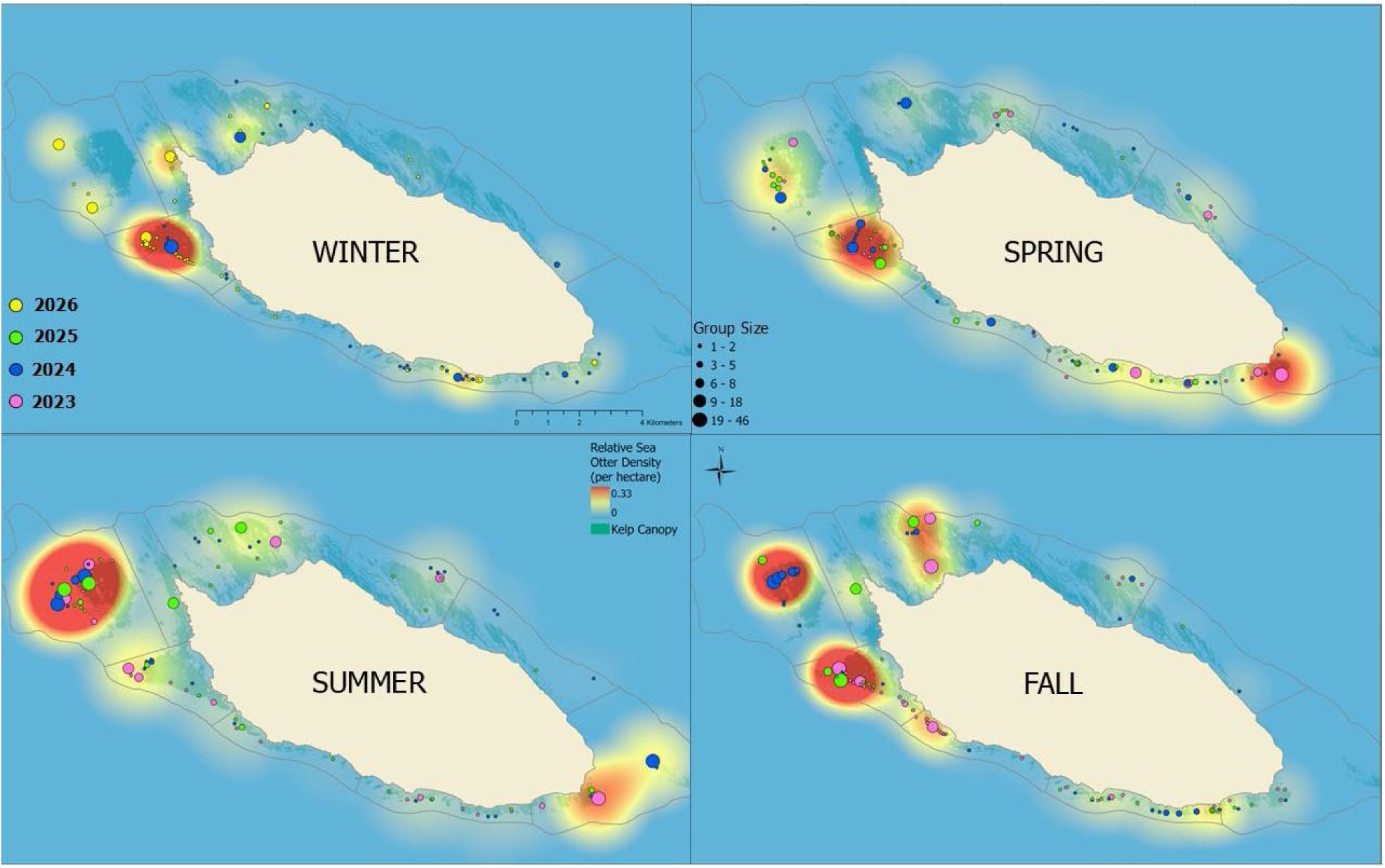
Seasonal distribution of relative sea otter density during surveys completed annually in winter (January/February), spring (April/May), summer (July/August), and fall (December) from 2023 to 2026, San Nicolas Island, California. Sea otter locations were aggregated across surveys for each season to develop kernel density surfaces representing the relative density of total counts distributed around San Nicolas Island, California. The legends for group size in the spring quadrant and relative sea otter density in the summer quadrant apply to all seasons. Mapped kelp canopies (shaded dim green) are derived from aerial imagery merged from the fall seasons of 2017-2019 and displayed for reference; however, actual canopy cover at the time of sea otter surveys can vary due to seasonal expansion and contractions caused by storm activity and other factors.

To estimate and test seasonal differences in the spatial distribution of sea otters, we aggregated sea otter counts within the west end, north side, east end, and south side sections (fig. 1). For surveys in the recent period from 2023 through February 2026, we detected significant differences in distribution among sections between years (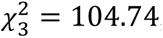, *p* < 0.001), seasons (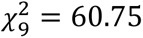, *p* < 0.001), and combinations of years and seasons (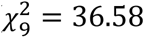, *p* < 0.001). We also detected significant differences in distribution among sections during the three-year periods of 2017–2019, 2020–2022, and 2023 to winter 2026 (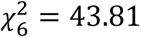, *p* < 0.001), with each period differing pairwise from one another (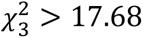, *p* < 0.001). Figure 9 shows the distribution of sea otters in these four sections over time.

**Figure 9.**
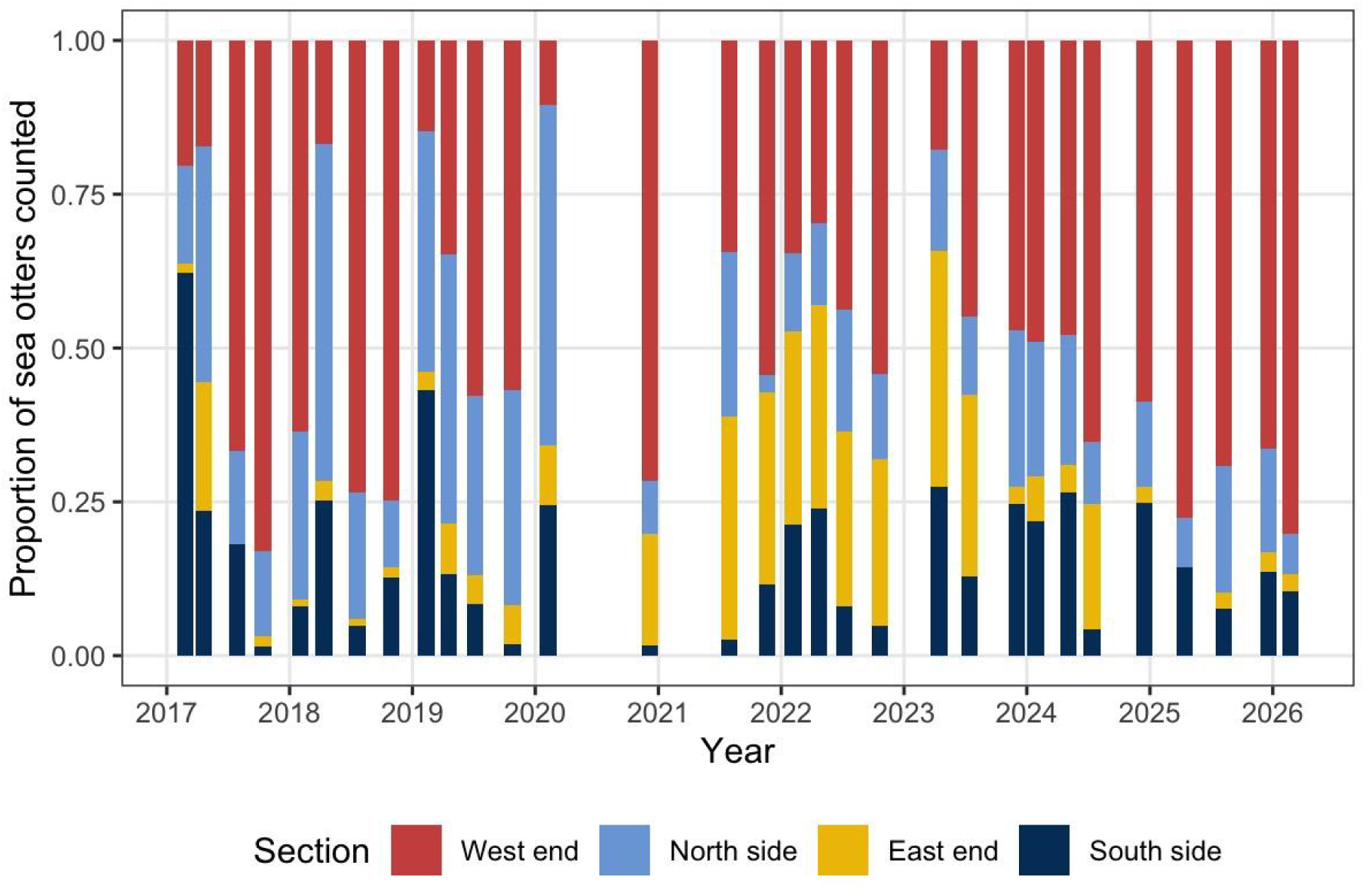
Proportional distribution of sea otters within four sections (West end = Areas 1-3, North side = Areas 4-6, East end = Area 7, South side = Areas 8-9) around San Nicolas Island, California, observed during seasonal surveys (winter, spring, summer and fall) from winter 2017 to winter 2026.

### Foraging

Foraging data were collected opportunistically as time and viewing conditions allowed. Consequently, the number of foraging observations varied by survey and location. A total of 32 foraging bouts were observed from 2023–winter 2026, bringing the total number of sea otter foraging bouts recorded since 2017 to 252. A total of 461 foraging dives were recorded between 2023–winter 2026, bringing the total number of foraging dives observed since 2017 to 3,701 (tables 3, 4). From 2023–2026, most foraging bouts were recorded on the east end (43.8 percent) and when combined with the west end (37.5 percent), the two ends of the island, which are the most accessible, make up 81.3 percent of all the foraging bouts recorded (table 3). Of the 461 total foraging dives recorded from 2023–winter 2026, more dives were recorded on females (332) than males (29) (table 4).

**Table 3.** Number of sea otter foraging bouts observed by season, sex, age class, and section of San Nicolas Island, California, 2023–2026, summarized by year and for all years combined. This report includes only the first (winter) survey for 2026. Dashes (-) indicate data not yet available.

| Sea otter foraging bouts | Year |  |  |  | All<br>year<br>s |
| --- | --- | --- | --- | --- | --- |
|  | 2023 | 2024 | 2025 | 2026 |  |
| Total foraging bouts | 8 | 15 | 8 | 1 | 32 |
| Season |  |  |  |  |  |
| Winter | 0 | 10 | 0 | 1 | 11 |
| Spring | 4 | 3 | 3 | - | 10 |
| Summer | 2 | 0 | 1 | - | 3 |
| Fall | 2 | 2 | 4 | - | 8 |
| Sex |  |  |  |  |  |
| Females | 5 | 10 | 6 | 1 | 22 |
| Males | 1 | 0 | 1 | 0 | 2 |
| Unknown sex | 2 | 5 | 1 | 0 | 8 |
| Age class |  |  |  |  |  |
| Adults | 6 | 10 | 6 | 1 | 23 |
| Aged Adults | 0 | 0 | 0 | 0 | 0 |
| Sub-adults | 2 | 2 | 2 | 0 | 6 |
| Juveniles | 0 | 0 | 0 | 0 | 0 |
| Unknown age | 0 | 3 | 0 | 0 | 3 |
| Section |  |  |  |  |  |
| West side | 0 | 6 | 5 | 1 | 12 |
| North side | 1 | 3 | 0 | 0 | 4 |
| East side | 6 | 6 | 2 | 0 | 14 |
| South side | 1 | 0 | 1 | 0 | 2 |

**Table 4.** Number of sea otter foraging dives observed by season, sex, and diving outcomes at San Nicolas Island, California, 2023–February 2026, summarized by year and for all years combined.

| Sea otter foraging dives | Year |  |  |  | All years |
| --- | --- | --- | --- | --- | --- |
|  | 2023 | 2024 | 2025 | 2026 |  |
| Foraging dives | 140 | 238 | 78 | 5 | 461 |
| Season |  |  |  |  |  |
| Winter | 0 | 145 | 0 | 5 | 150 |
| Spring | 84 | 41 | 26 | 0 | 151 |
| Summer | 21 | 0 | 9 | 0 | 30 |
| Fall | 35 | 52 | 43 | 0 | 130 |
| Sex |  |  |  |  |  |
| Females | 98 | 165 | 64 | 5 | 332 |
| Males | 20 | 0 | 9 | 0 | 29 |
| Unknown sex | 22 | 73 | 5 | 0 | 100 |
| Outcomes |  |  |  |  |  |
| Successful dives | 108 | 140 | 52 | 5 | 305 |
| Unsuccessful dives | 29 | 59 | 24 | 0 | 112 |
| Other | 1 | 33 | 1 | 0 | 35 |
| Unknown outcomes | 2 | 4 | 1 | 0 | 7 |
| Known outcomes | 138 | 234 | 77 | 5 | 454 |
| Known success or unsuccess | 137 | 199 | 76 | 5 | 417 |
| Percentage success rate (successes/dives) | 78.8 | 70.4 | 68.4 | 100 | 73.1 |
| Mean dives per bout | 17.5 | 15.9 | 9.8 | 5.0 | 14.4 |
| Mean diving duration (seconds) | 65.3 | 55.0 | 52.4 | 72.5 | 57.9 |
| Mean surface interval (seconds) | 42.5 | 57.6 | 55.0 | 74.4 | 52.8 |

We classified identifiable prey species into taxonomic groups based on highest prevalence in the diet (urchins, snails, bivalves, crabs) or high energetic content of less prevalent prey species (abalone, lobsters, octopus) and lumped all other species (other). We used Bayesian Monte-Carlo simulations to estimate the likely caloric values of unidentified prey based on partial data about the prey, such as their size and the lowest taxonomic level to which they could be identified. These simulations use empirical information about the likelihood of different prey species and the associated caloric values for their size class, based on the frequency with which specific species and sizes were identified in the data. This process enabled us to derive 95- percent credible intervals to account for prey uncertainty. These simulations require robust sample sizes to provide sufficient information on prey. Because our foraging observations in the 2020–2026 period were fewer compared to those of previous years, we aggregated those data across years to calculate an average energy intake rate for the 2020–2026 period. Additionally, we calculated energy intake rates separately by year from 2017 to 2019 and for the combined 2003–2006 period.

From 2023-2026, sea urchins, including the red sea urchin (*Mesocentrotus franciscanus*) and purple sea urchin (*Strongylocentrotus purpuratus*), composed the slight majority (52.1 percent) of identified prey types among all successful diving outcomes, with the remaining successful identified diving outcomes comprised mostly of snails (20.2 percent), bivalves (17.2 percent), and crabs (9.7 percent; Table 5). Twenty-seven percent of successful dives (87 of 325) observed during 2023-2026 resulted in unidentified prey items (table 5). Overall urchin consumption decreased from the prior three-year period (2020-2022) when urchins made up two thirds of the identified prey items. The one third decrease in urchin consumption was balanced by a doubling of snail, bivalve, and crab consumption (fig. 10).

**Figure 10.**
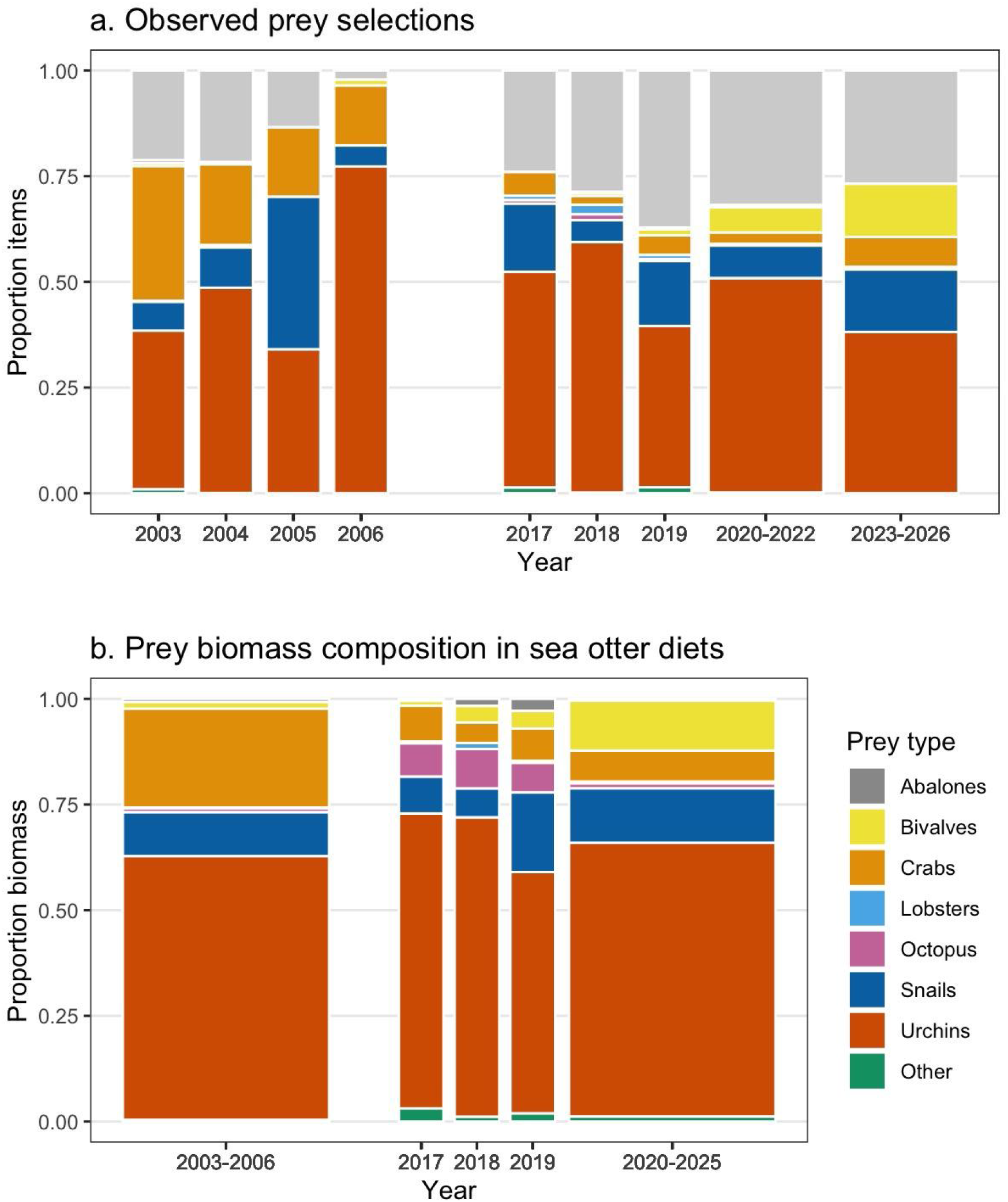
Proportions of a) observed prey types, and b) prey biomass composition in sea otter diets during foraging surveys at San Nicolas Island, California, 2003–2006 and 2017–2026. There were too few data in winter 2026 to include with biomass composition, and some years were aggregated due to smaller sample sizes and displayed in wider bars to represent multiple years.

**Table 5.**
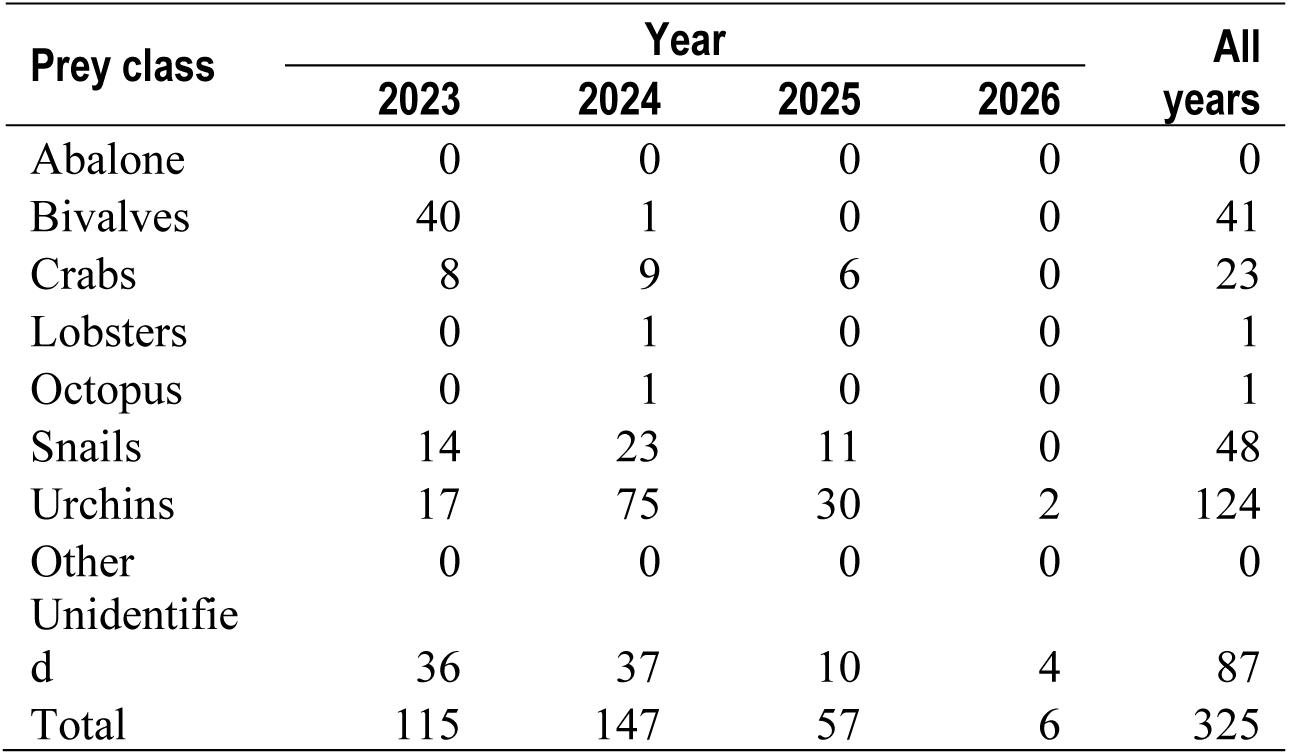
Number of prey categorized into eight prey classes observed during foraging surveys at San Nicolas Island, California, 2023–February 2026.

| Prey class | Year |  |  |  | All years |
| --- | --- | --- | --- | --- | --- |
|  | 2023 | 2024 | 2025 | 2026 |  |
| Abalone | 0 | 0 | 0 | 0 | 0 |
| Bivalves | 40 | 1 | 0 | 0 | 41 |
| Crabs | 8 | 9 | 6 | 0 | 23 |
| Lobsters | 0 | 1 | 0 | 0 | 1 |
| Octopus | 0 | 1 | 0 | 0 | 1 |
| Snails | 14 | 23 | 11 | 0 | 48 |
| Urchins | 17 | 75 | 30 | 2 | 124 |
| Other | 0 | 0 | 0 | 0 | 0 |
| Unidentified | 36 | 37 | 10 | 4 | 87 |
| Total | 115 | 147 | 57 | 6 | 325 |

We examined the composition of sea urchin species, which was the most identified prey among successful sea otter foraging dives. Red sea urchins generally are larger (median diameter = 53 mm, interquartile range [IQR; 25–75-percentile] = 41–65 mm) at SNI and thus have higher caloric value than purple sea urchins (median diameter = 28 mm, IQR = 21–36 mm; Kenner and Tomoleoni, 2020). Because sea otters are size-selective predators, they tend to prefer red sea urchins when larger size classes of red sea urchins are more available. As such, changes in the ratio of red to purple sea urchins consumed can correlate with prey availability. We inspected this ratio during the 2020–2022 and 2023–2026 periods and compared it to ratios calculated annually from 2003 to 2006 and 2017 to 2019 when similar sea otter foraging surveys were done at SNI. Results indicated a decreasing trend in the ratio of red to purple sea urchins consumed, detectable in 2017–2019 and pronounced by 2020 and later (fig. 11).

**Figure 11.**
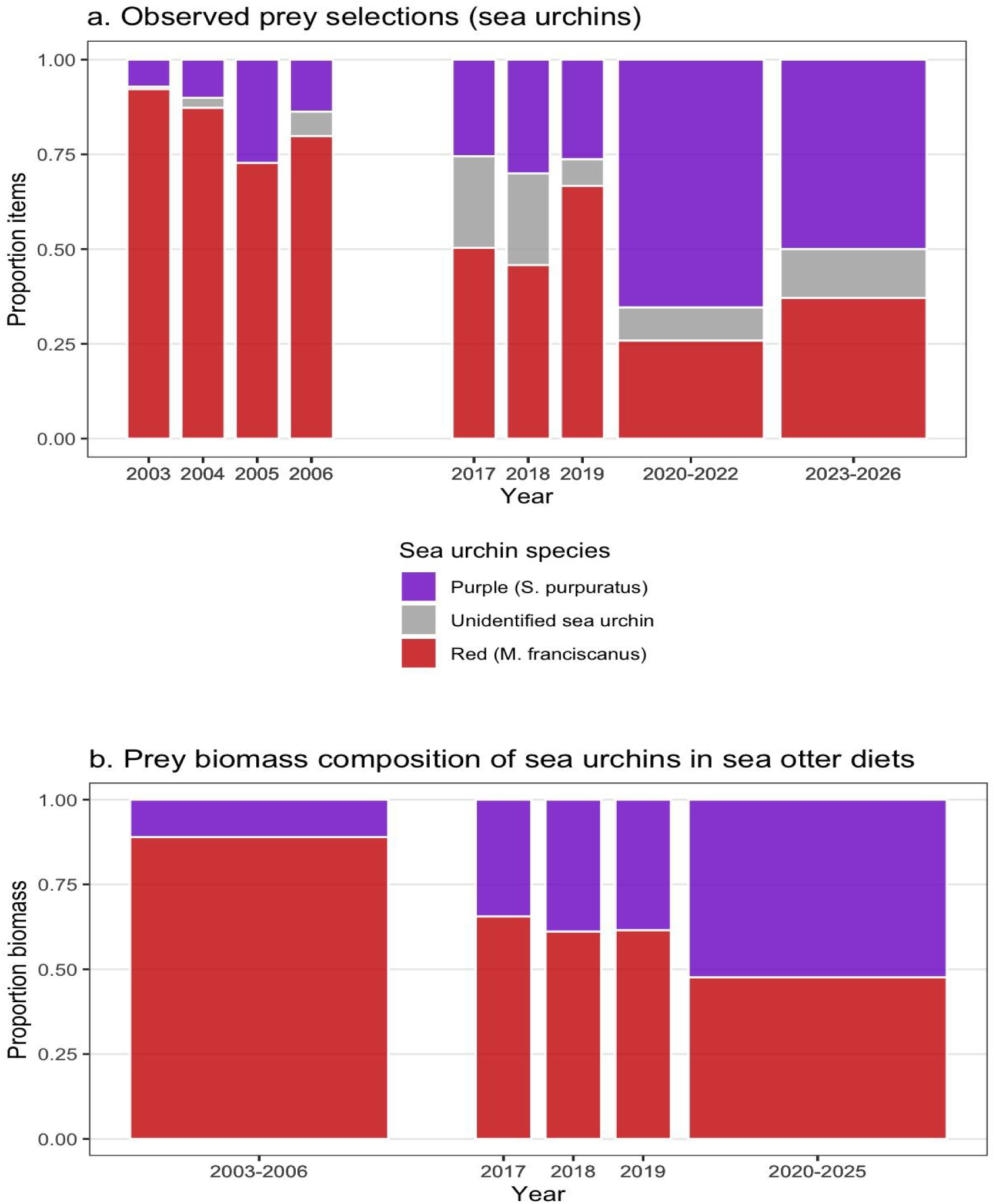
Proportions of a) observed sea urchin prey species (*Mesocentrotus* and *Strongylocentrotus*), and b) sea urchin biomass composition in sea otter diets during foraging surveys at San Nicolas Island, California, 2003–2006 and 2017–2026. There were too few data in winter 2026 to include with biomass composition, and some years were aggregated due to smaller sample sizes and displayed in wider bars to represent multiple years.

Simulated estimates of energy intake rates during 2020–2025 averaged 7.7 kcal/min (95- percent CI = 6.6–9.1 kcal/min), which continued a nearly linear pattern of decrease that started in 2017 (9.0 kcal/min; 95-percent CI = 7.6–10.7 kcal/min). Despite the consistency of this pattern going through the intervening years, 2018 and 2019, the credible intervals of these estimates overlap, suggesting the decrease is statistically significant. The estimated energy intake rate from 2017 was close to that of the previous 2003–2006 surveys (8.9 kcal/min; 95-percent CI = 8.0–9.9 kcal/min) (fig. 12). Dietary diversity calculated from simulated estimates of prey class biomass trended positively over time (Shannon-Wiener index [SWI] = 1.26, 1.53, 1.58, and 1.73 for the period 2003-2006 and years 2017, 2018, and 2019, respectively), but this pattern of increase did not continue into 2020–2026 (SWI = 1.59).

**Figure 12.**
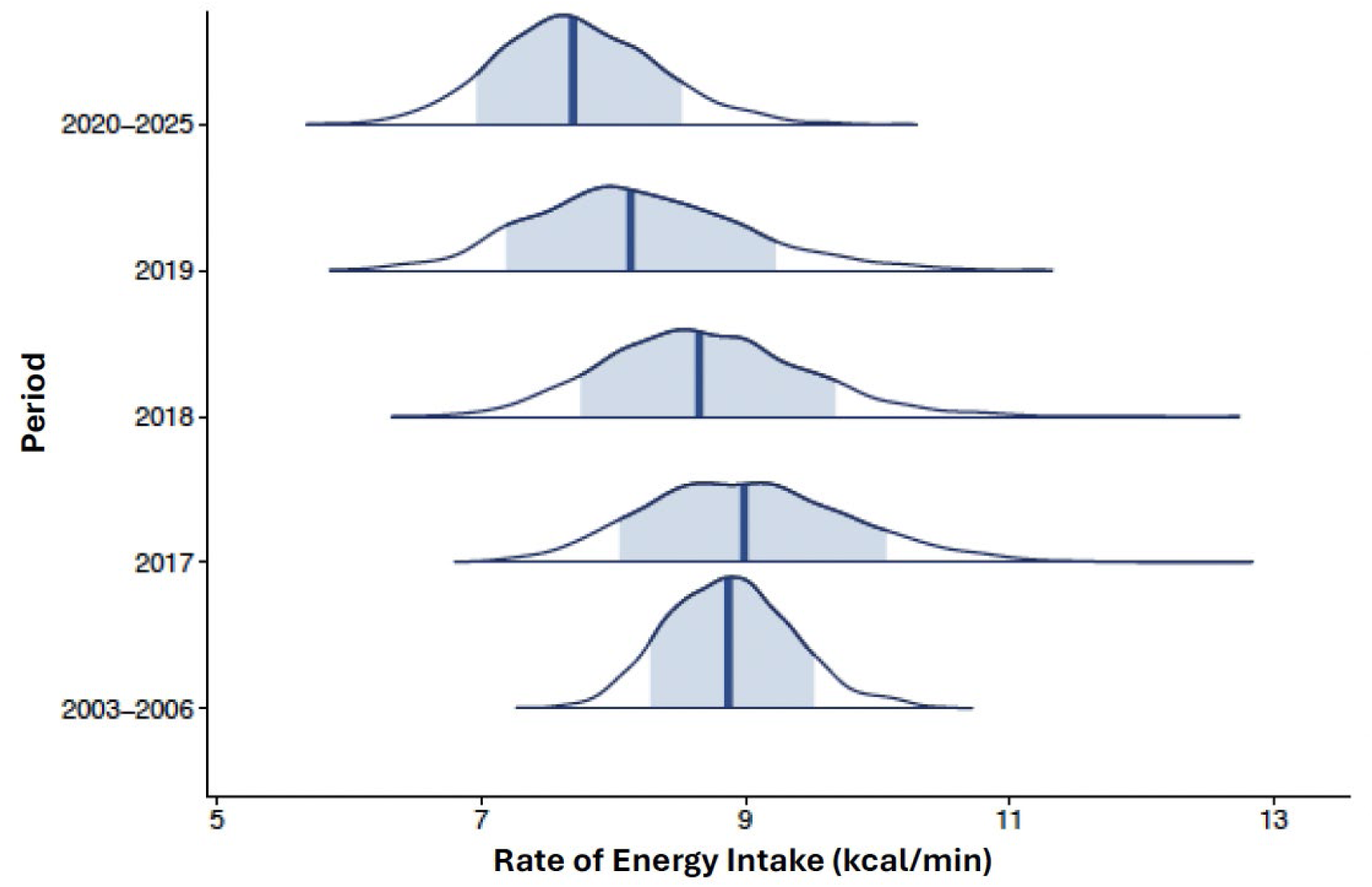
Rates of energy intake (kilocalories per minute) estimated for sea otters observed during forage surveys at San Nicolas Island, California, between 2003 and 2025. Estimated rates are presented as posterior probability distributions with median estimate (vertical line) and 95% credible intervals (shaded values) for combined years in 2003–2006, individual years in 2017–2019, and combined years in 2020– 2025. There were too few data in winter 2026 to include with estimates of energy intake rates.

## Discussion

Our surveys reveal the dynamic nature of the sea otter population at SNI. Island-wide counts vary between surveys, with occasional periods of decline and more frequent periods of growth (figs. 2 & 4). Overall, the population has sustained a positive growth trajectory (7.4 percent per year; 95-percent CI = 6.5 to 8.4 percent per year) since 1990. However, since monitoring under the Southern Sea Otter Military Readiness Area Monitoring and Research Plan began in 2017, we have observed population growth rates taper, from 19.6 percent per year (95- percent CI = 9.9 to 30.1 percent per year) from 2017 through 2020, to 5.5 percent per year (95-percent CI = –8.2 to 14.2 percent per year) from 2020 through 2023, to –8.1 percent per year (95- percent CI = –1.1 to –14.6 percent per year) from 2023 through February 2026. The recent surveys mark the first time we have observed a statistically significant decline (8.1 percent) at SNI since the earliest years of post-translocation monitoring (fig. 4). Subsequent surveys will reveal whether this decline is temporary or part of a longer-term trend. Depending on the results of future surveys, a continuing decline could eventually trigger tier 3 monitoring according to the Southern Sea Otter Military Readiness Area Monitoring and Research Plan.

The count from the most recent sea otter survey in February 2026 was 106 total sea otters (table 2), 40 sea otters fewer than the final survey included in the previous (2020-2023) report (Yee and others, 2023). The biggest driver of the recent population decline at SNI appears to be the sudden disappearance of the east end group of more than 40 sea otters in the fall of 2023. The data indicate that the east end sea otters did not simply shift to using the west end of the island, because the island-wide count also declined significantly when the east end raft disappeared. This strongly suggests that most of the east end sea otters either left the island or died, with the former being the more likely explanation. In June 2024, less than a year after the disappearance of the east end group, a single male sea otter was observed foraging and resting off La Jolla in San Diego County. This area is well beyond the southern range boundary (near Gaviota in Santa Barbara County) for sea otters on the California mainland, and there is a very strong likelihood that this sea otter came from SNI. We also hypothesize that the large east end group at SNI was likely a male group, as pups were typically not observed in this group during our surveys. Large male sea otter groups also form among the California mainland sea otter population and often exhibit interesting or atypical behavior. These male groups can persist for years, but the animals eventually disperse, likely in search of mating opportunities. We hypothesize that the large east end raft was a male group that dispersed, leaving the island and triggering the recent overall population decline at SNI. If our hypothesis is correct, then we would expect a period of stabilization in the island-wide counts at these lower overall numbers compared to the previous 3-year period, followed by a return to positive population growth. If the annual rate of population growth continues along a negative trend, then there would be reason to suspect that something other than the emigration of the east end group could be driving the decline.

The distribution of otters at SNI has shifted once again as compared to the 2020-2023 report, this time returning to pre-2017 geographic patterns characterized by heavier use of the west end, lighter use of the south side, and very low use of the east end and north side (fig. 5, 6). The island-wide distribution was also characterized by seasonal shifts, with peak east end occupation occurring in spring and summer; however, once the east end group departed post-summer 2023, the densities of sea otters counted at the east end remained very low in all seasons. One of the areas most commonly occupied by sea otters, The Boilers (Area 3), showed a 3- season usage pattern, with the highest densities in summer and fall when kelp canopy extent was at its normal maximum. In stark contrast, there were no sea otters detected at The Boilers in the winter season for 2024 and it was noted that no kelp was present during that survey. Our most recent winter survey (2026) detected both significant sea otter numbers and significant kelp canopy at The Boilers. Because sea otters frequently use persistent kelp canopy for resting, especially in open ocean conditions like those at SNI, and because kelp forests typically provide a greater prey base for foraging sea otters, seasonal changes in kelp are likely to be an important driver of the seasonal variations in sea otter distributions that we observed.

Energy intake rates of sea otters is one indicator of population status. Foraging habits are known to change as sea otter populations increase and food resources become more difficult to obtain, and these density-dependent processes can signal that a population is approaching carrying capacity (Estes and others 2003; Tinker and others 2008; Tinker and others, 2012; Tinker and others, 2019). There were indications that energy intake from foraging activity might have slightly decreased annually from 2017 to 2019, with the decreasing trend continuing into 2020–2025. However, fewer samples of foraging observations collected annually during the latter period precluded us from calculating annual estimates or a precise trend in energy intake. Instead, we aggregated the recent data into multi-year periods, and although there was a temporal pattern of decline in the energy intake rate, the credible intervals for those estimates widely overlapped, and we could not conclude a statistical change. Additional data collection could have allowed higher temporal resolution and stronger interpretations. A decline in forage intake rate in conjunction with an increase in population size would be expected for a population that is beginning to experience density-dependent effects.

The opportunistic nature of foraging data collection means that there is some location associated bias inherent to foraging data. More foraging data are collected where higher densities of sea otters are found. Also, more foraging data are collected from areas where sea otters are closer to shore and where good vantage points are easily accessible. More remote areas of the island requiring hours of hiking or locations with lower sea otter densities will always have sparser foraging data compared to higher density and more easily accessed areas such as Area 1 in the west end. The arrival of a group of 40+ sea otters at the east end (Area 7), another accessible location where sea otters occur close to shore, created an opportunity to augment foraging data (802 dives between 2022–2024). When this group disappeared, our total foraging data numbers declined (83 dives from 2025–2026) as our locations for collecting foraging data became more limited and less ideal. These logistical biases could mean that our estimates do not fully represent all sea otters at San Nicolas Island; however, the extent to which these biases could impact our estimates is unknown.

Foraging results should be interpreted cautiously, as the nature of foraging surveys can introduce potential biases that do not occur with population surveys. Survey effort at SNI was prioritized toward counting sea otters, which occurred on days with the most optimal viewing conditions within each survey interval. Sea otters were systematically counted around the island as synchronously as possible, always within a day, to minimize counting errors due to sea otter movements. These surveys were repeated over multiple days to minimize undercounting due to fluctuating survey conditions, sea otter grouping, behavior and detectability, and we recorded the highest single-day count for each survey as a robust estimate of the population. In contrast, foraging surveys were done opportunistically as time and viewing conditions allowed. As a result, the amount of available foraging information is limited and uneven across surveys, locations, and individual sea otters. Because of limited observations, we aggregated foraging data to compute and compare energy intake rates by year, season, and sex. However, if foraging activity differs by any of these factors, then those effects can potentially confound with these unbalanced aggregations when comparing other factors. Systematic biases also could exist. Foraging observations have an unavoidable bias toward prey acquired during daytime. Previous studies have shown that sea otter foraging bouts occur both during day and night with equal frequency (Tinker and others, 2006). Additionally, data from time-depth recorders implanted in the 2003–2005 SNI study show variation between daytime and nighttime dive characteristics that could result from sea otters seeking different prey nocturnally (Newsome and others, 2010). Certain high-value mobile invertebrate prey species like lobster, crabs, and octopus are more active at night, and therefore could be more available to nocturnally foraging sea otters. Furthermore, foraging observations require translation into energy intake values based on prey species and size, and unidentified items are translated into an average value for the size class. The accuracy of these translations depends on detailed and reliable information on the nutritional content of prey items, which have been documented extensively by Oftedal and others (2007), although it remains a work in progress to update and expand these keys for different locations and time periods.

Density-dependent processes, such as changes in sea otter foraging habits associated with changes in densities of sea otters, are known to occur for growing populations of sea otters as they deplete preferred prey resources and must increasingly compete for prey, generally indicated by reduced energy intake rates and increased dietary diversity. We did not find significant changes in the energy intake rates of foraging sea otters nor a clear increase in dietary diversity, but there were other signs that foraging patterns have potentially begun to change. We observed a pronounced trend from 2003 to 2026 of sea otters increasingly foraging on purple sea urchins instead of the preferred larger red sea urchins, as well as a proportionally higher consumption of bivalves in recent years. Annual subtidal surveys contained evidence supporting an overall increase in purple sea urchins on the island’s east end (Kenner and Tomoleoni, 2021) where a large group of (presumably male) sea otters took up residence from 2021–2023 (Yee and others, 2023). We also estimated slight declines in energy intake rates from 2017–2019 and continuing into 2020–2025, but the differences are small, and with small sample sizes and wide confidence intervals we are unable to assess change. A continuation of these monitoring efforts and an increase in samples can provide further insights into the spatial and foraging dynamics of the population of sea otters at SNI and its relationship to the surrounding nearshore ecosystem.

The single biggest gap in our existing knowledge of the SNI sea otter population is their movement patterns, particularly immigration to and emigration from SNI. This report, which documents the sudden disappearance of the east end group, highlights the need for better data on the movement patterns of SNI otters. Traditionally, sea otter movement patterns are established by tagging and tracking studies, but this approach requires large-scale capture/tagging operations as well as an on-site tracking team locating sea otters daily using VHF radio telemetry. Additionally, while VHF telemetry allows researchers to locate sea otters within the bounds of San Nicolas Island, it does not easily allow for the detection of sea otters that leave the island. The USGS and colleagues have been pursuing new tagging technologies for sea otters that could potentially track sea otters anywhere they go, informing researchers and managers not only when they leave SNI and where they go once they leave, but also when they arrive at SNI and where they are coming from. Understanding sea otter movements will not only elucidate the location population dynamics at SNI but could reveal broader habitat use among the Channel Islands (especially San Clemente Island, which is included in the Southern Sea Otter Military Readiness Area Monitoring and Research Plan), mainland California, or even off Baja California, Mexico.

## Acknowledgments

This study was done in partnership with the U.S. Fish and Wildlife Service (USFWS) and the U.S. Navy (USN), with assistance from the California Department of Fish and Wildlife and the Monterey Bay Aquarium. We thank the USN for funding and facilitating this research project. J. Tomoleoni, J. Yee, and L. Bowen managed the project under the U.S. Geological Survey (USGS); E. Seacord facilitated access to San Nicolas Island and supported the project under USN, with coordination from L. Carswell under the USFWS; J. Tomoleoni, B. Hatfield, M. Staedler, and L. Carswell led multiple seasonal surveys, with E. Seacord, J. Fujii, G. Bentall, L. Konrad, C. Young providing collaborative support consistent with sea otter survey methodology developed by the USGS, USFWS, California Department of Fish and Wildlife, and Monterey Bay Aquarium; J. Tomoleoni performed GIS analysis; and T. Tinker, J. Yee, and L. Konrad performed data analysis, with T. Tinker providing expert guidance on sea otter forage analysis. All authors contributed to the writing of this report. We also thank numerous field observers throughout the 2023–2026 sea otter surveys at San Nicolas Island, California, including (in alphabetical order): Kevin Curran, Samantha Hamilton, Mike Harris, Traci Kendall, Karl Mayer, Keith Miles, Thomas Murphey, Teri Nicholson, Kate Riordan, and Jeff Wagner. We are grateful to John Ugoretz for contributing to the study design and the inception of this project, to James Estes who established the San Nicolas Island sea otter monitoring program, and to Claudia Makeyev and Greg Sanders for facilitating the continuation of this project over time. This report was vastly improved by comments and suggestions made by Josh Adams and Diane Elam. The data that support the findings of this report belong to the U.S. Navy and are not publicly posted, but are available from the corresponding author upon reasonable request and with explicit permission of the U.S. Navy. Any use of trade, firm, or product names is for descriptive purposes only and does not imply endorsement by the U.S. Government.

## Appendix 1. Figures of Pages from the Monitoring and Research Plan for Southern Sea Otter Military Readiness Area

**Figure 1.1.**
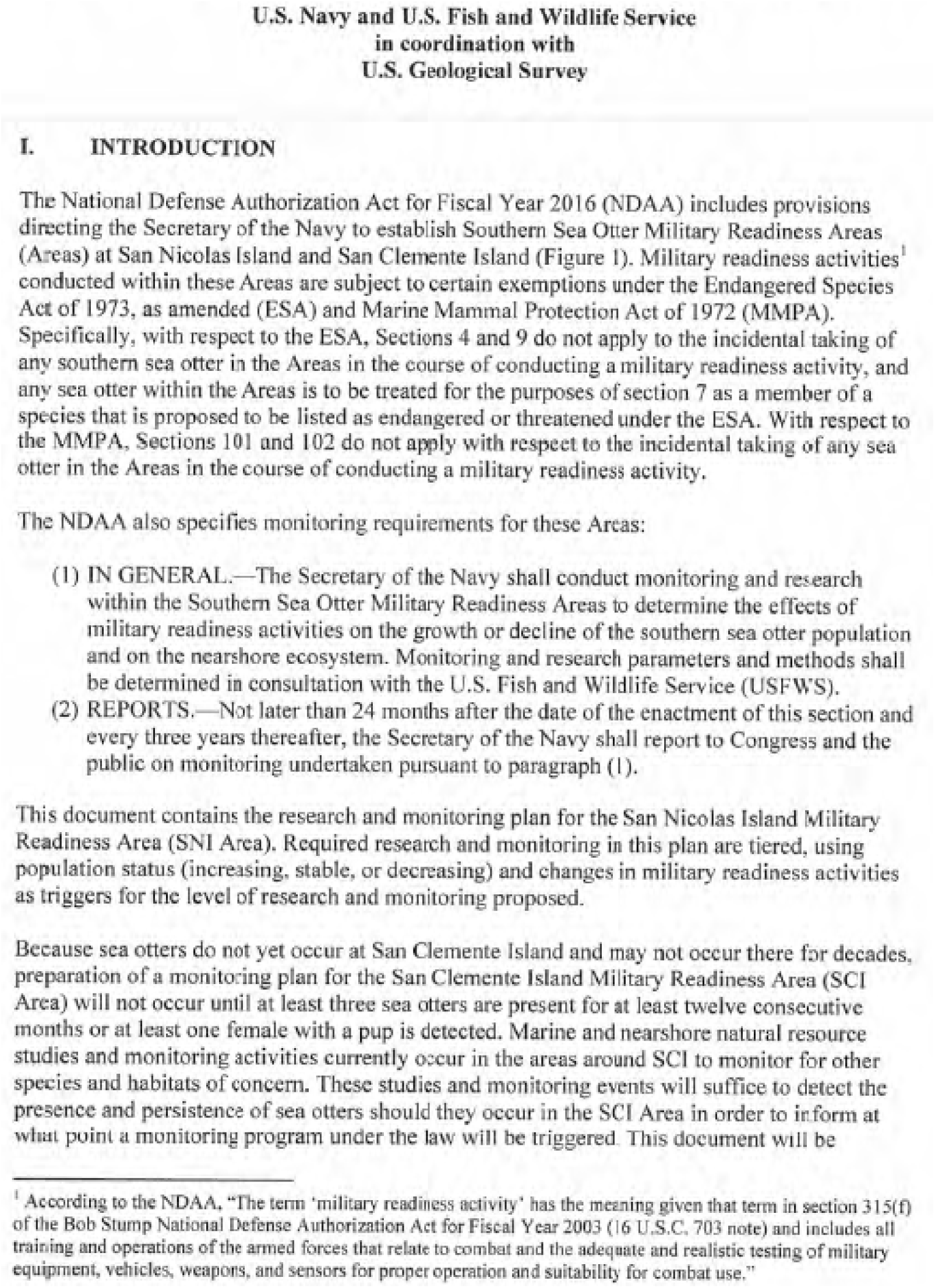
Page 1 of Monitoring and Research Plan for Southern Sea Otter Military Readiness Area.

**Figure 1.2.**
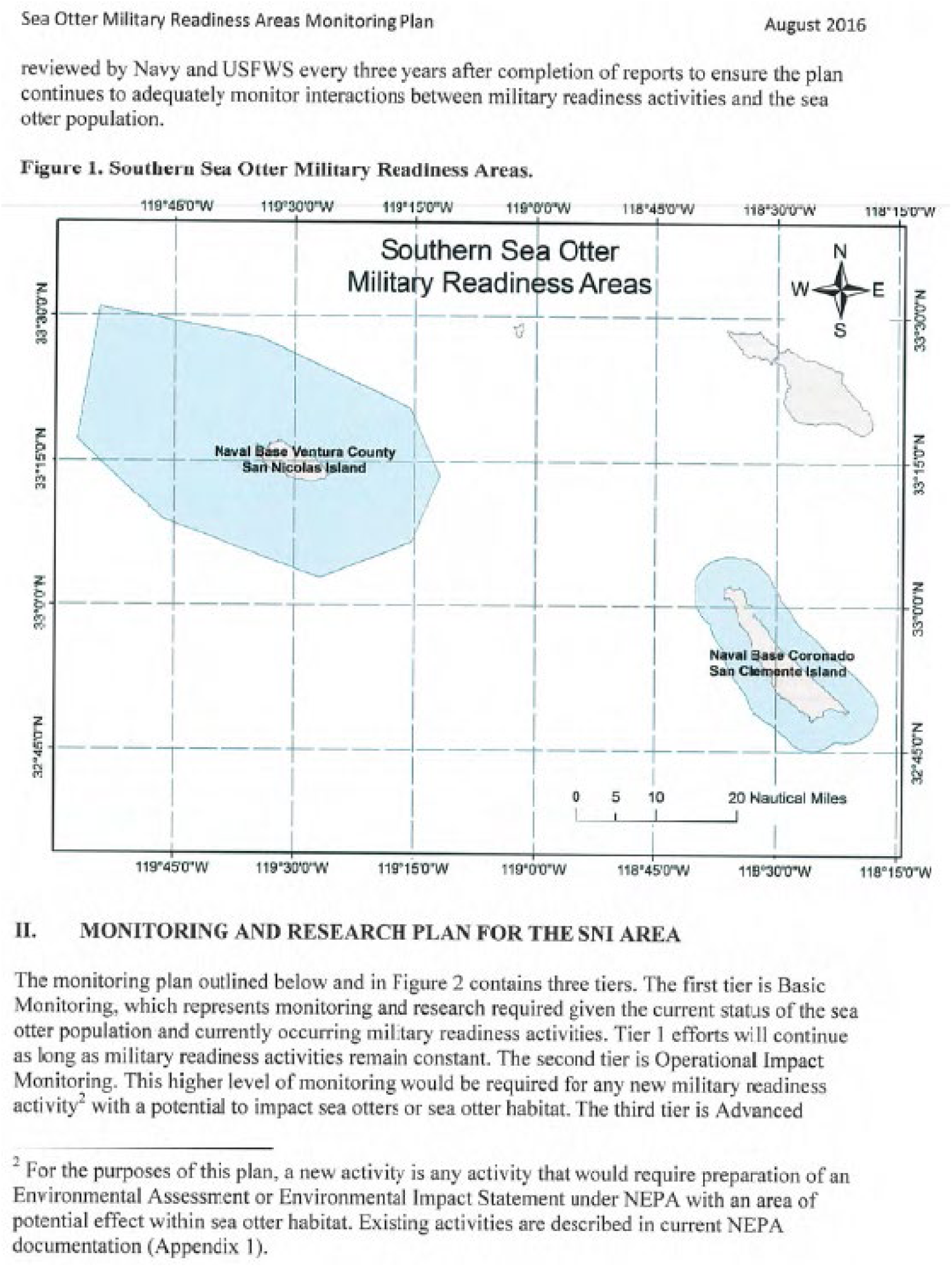
Page 2 of Monitoring and Research Plan for Southern Sea Otter Military Readiness Area.

**Figure 1.3.**
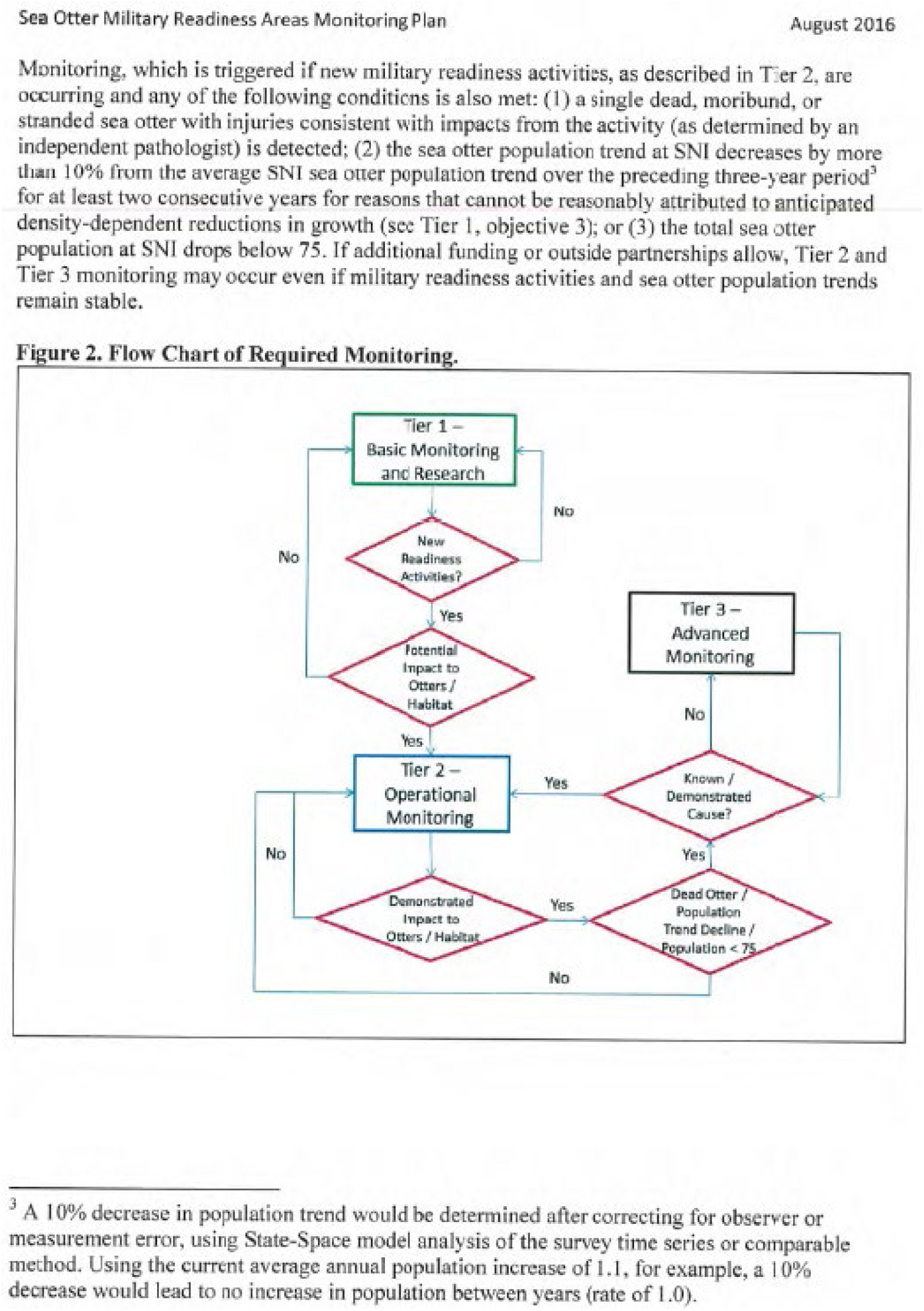
Page 3 of Monitoring and Research Plan for Southern Sea Otter Military Readiness Area.

**Figure 1.4.**
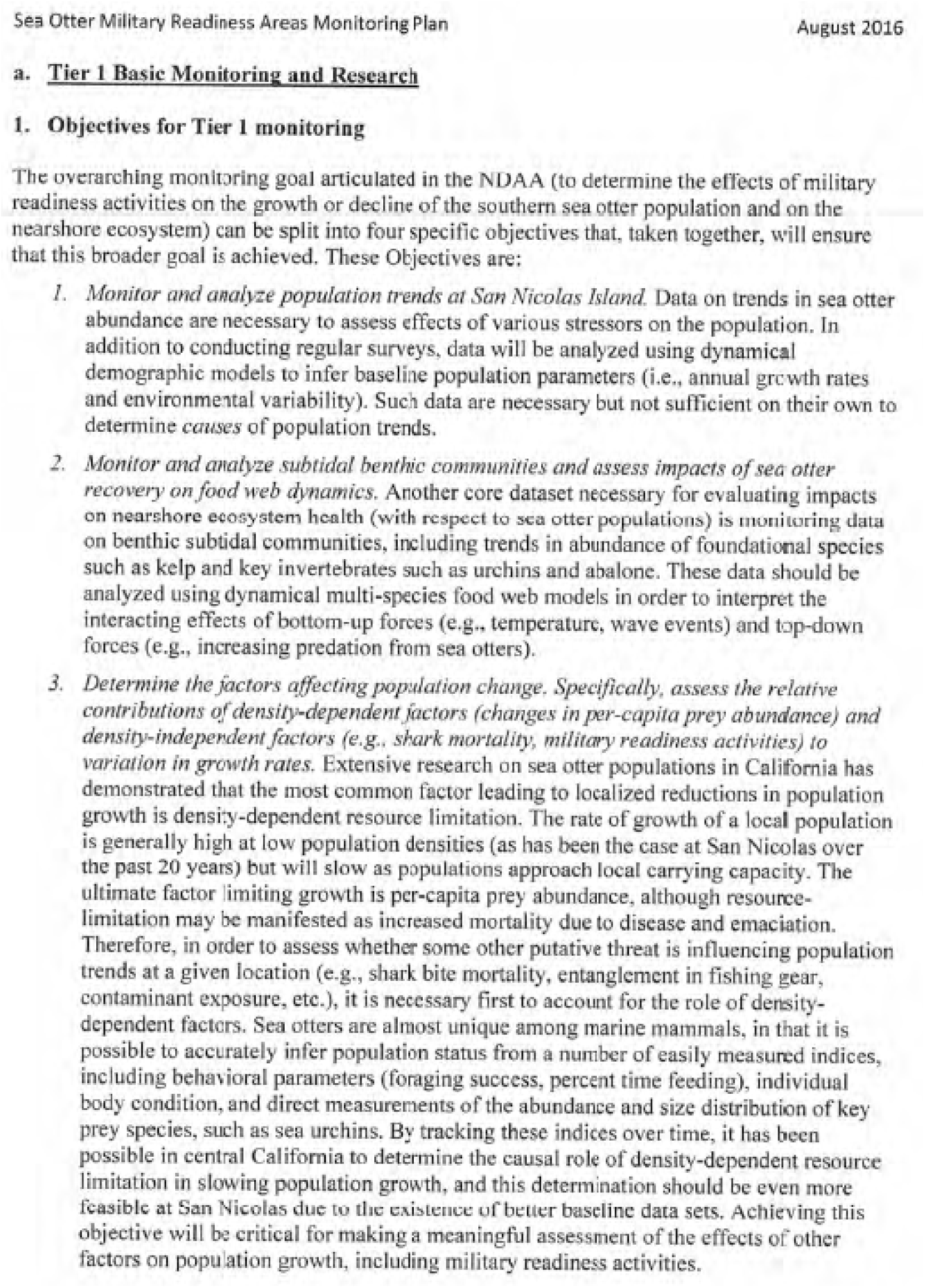
Page 4 of Monitoring and Research Plan for Southern Sea Otter Military Readiness Area.

**Figure 1.5.**
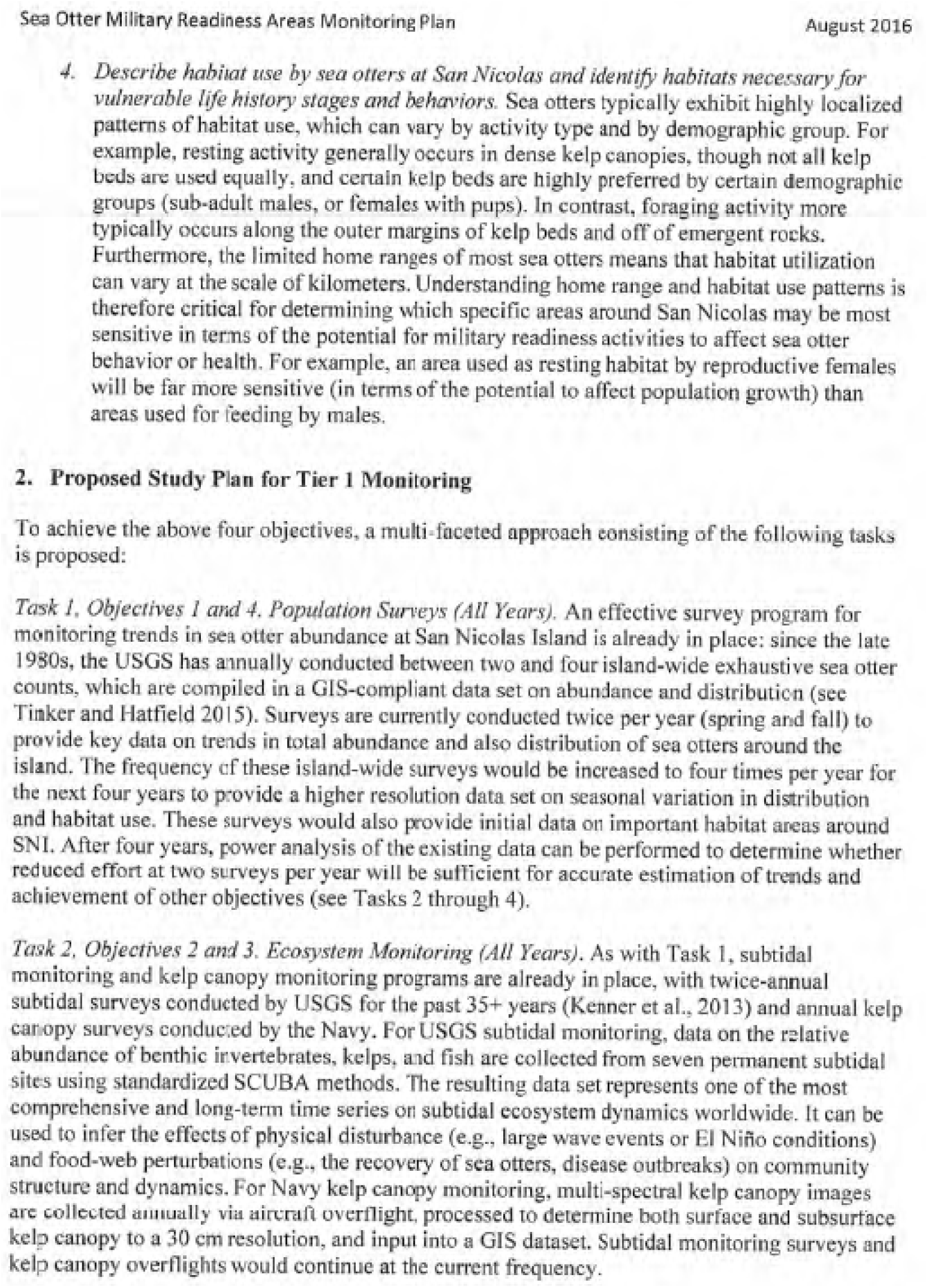
Page 5 of Monitoring and Research Plan for Southern Sea Otter Military Readiness Area.

**Figure 1.6.**
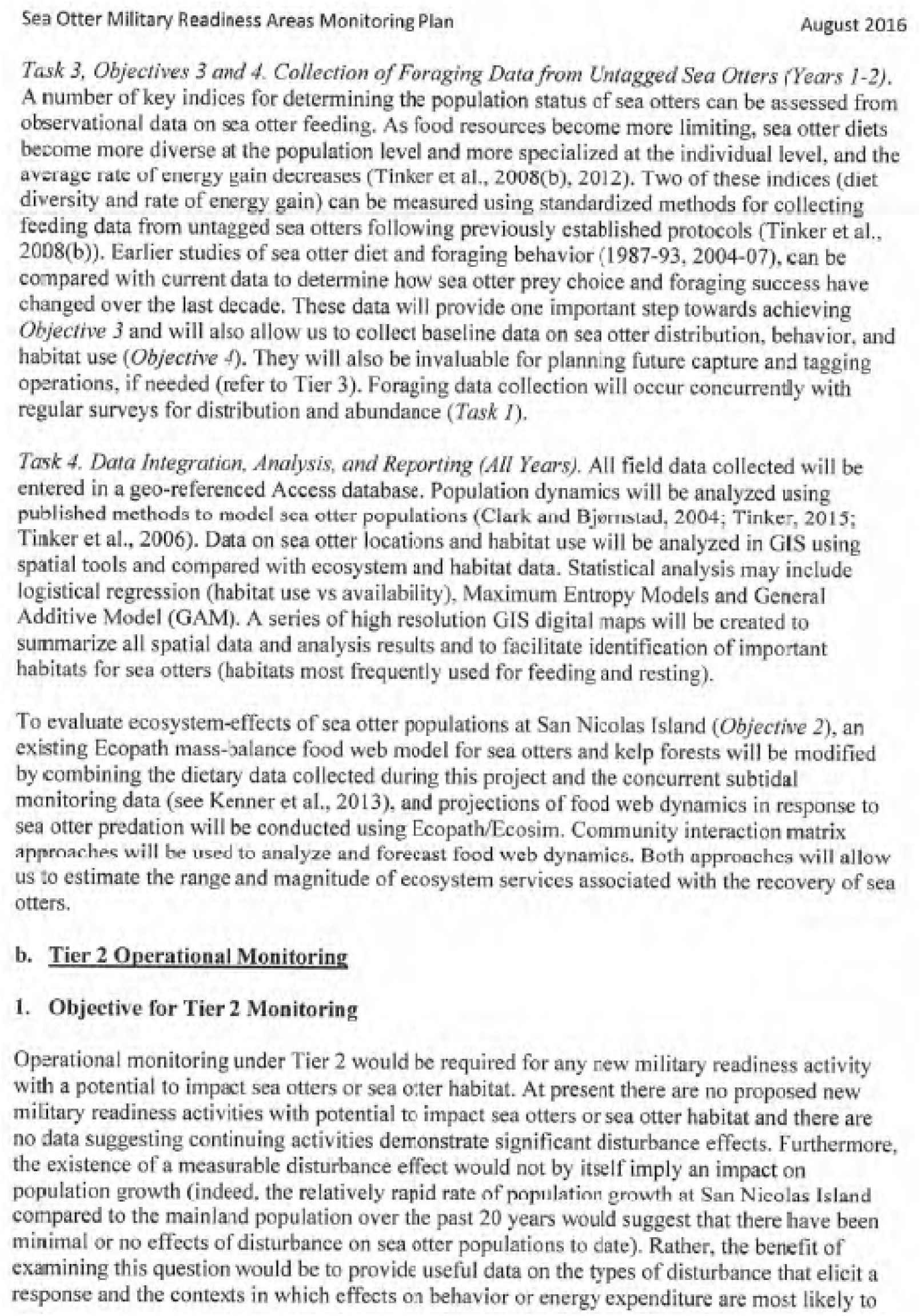
Page 6 of Monitoring and Research Plan for Southern Sea Otter Military Readiness Area.

**Figure 1.7.**
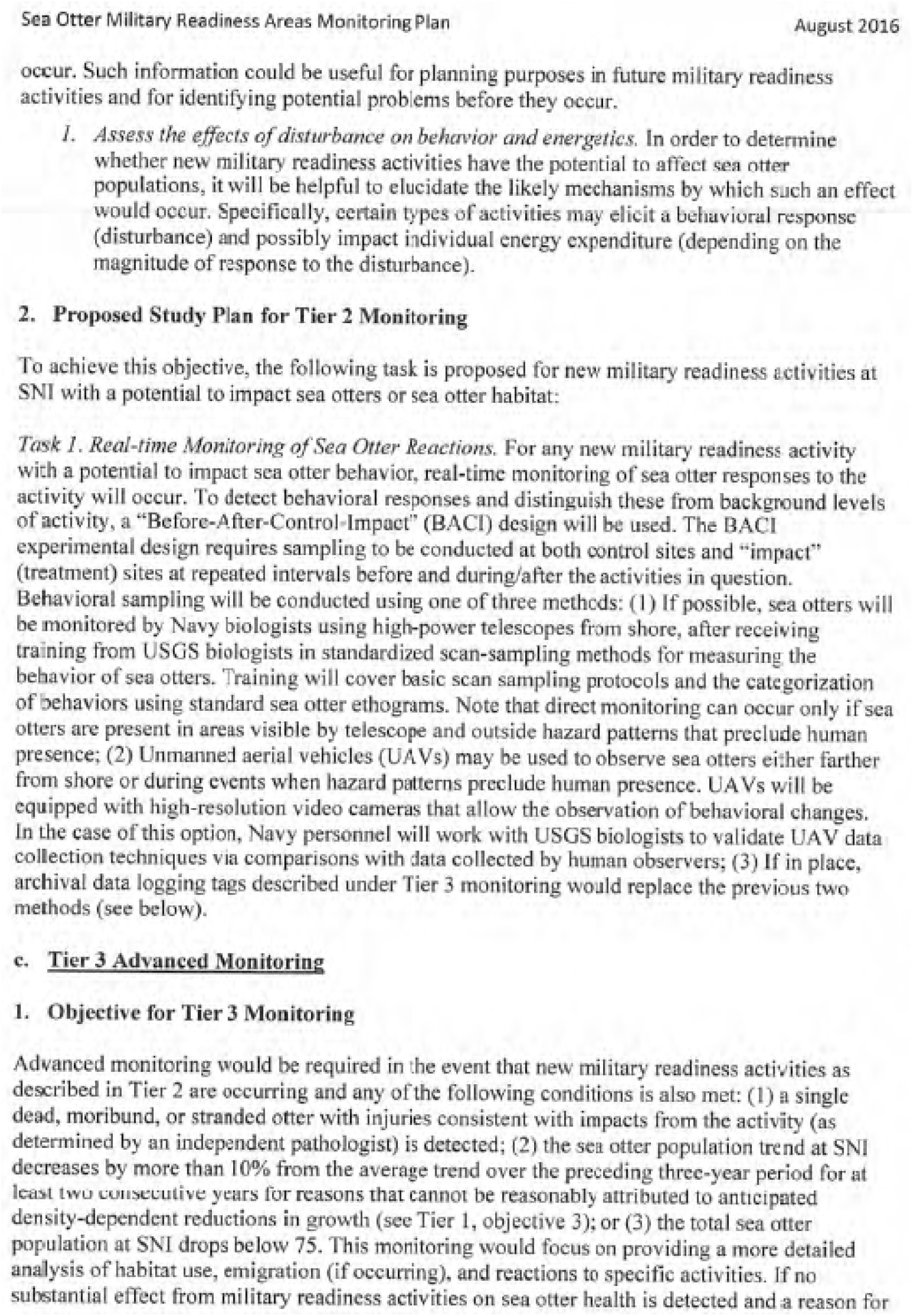
Page 7 of Monitoring and Research Plan for Southern Sea Otter Military Readiness Area.

**Figure 1.8.**
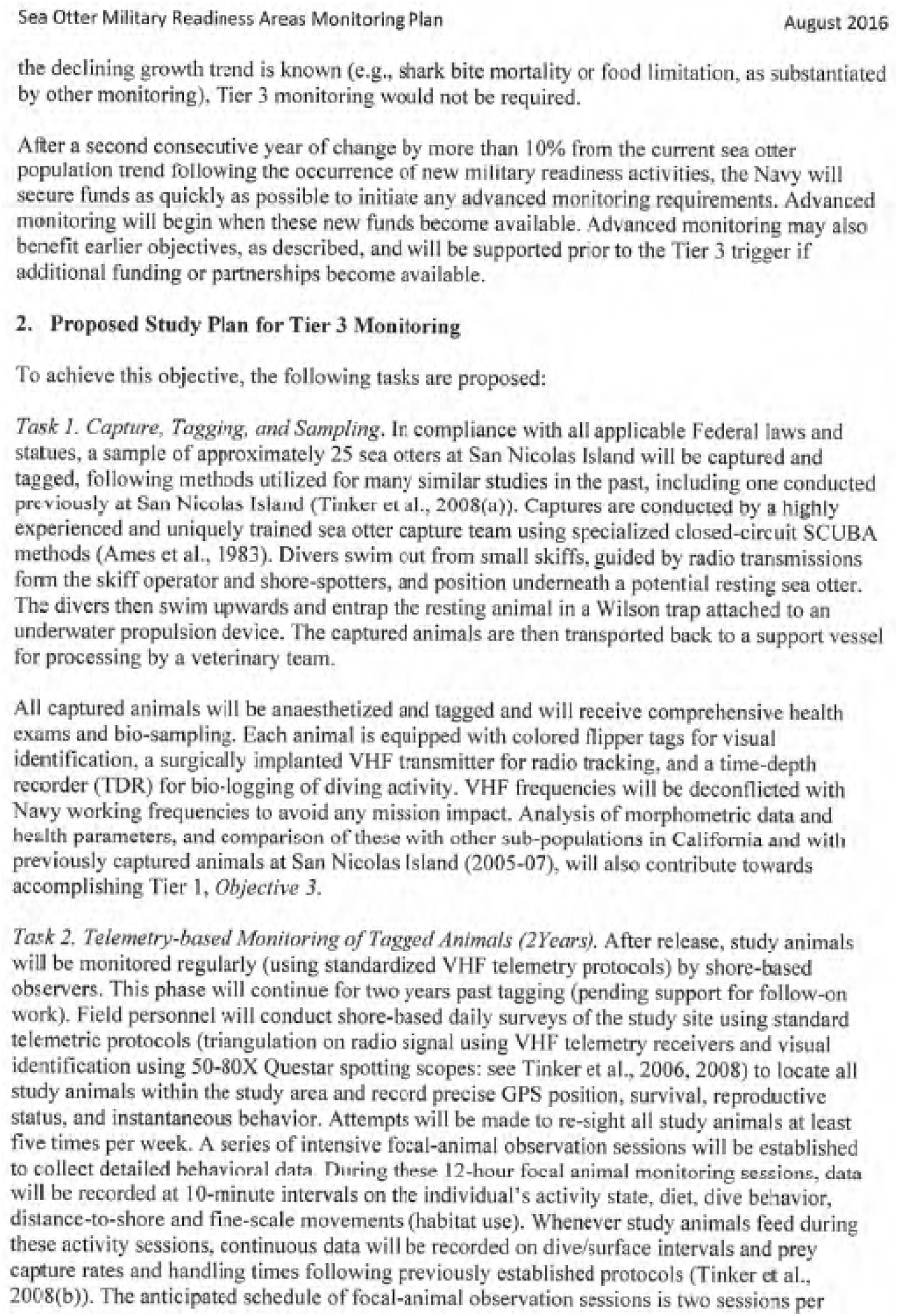
Page 8 of Monitoring and Research Plan for Southern Sea Otter Military Readiness Area.

**Figure 1.9.**
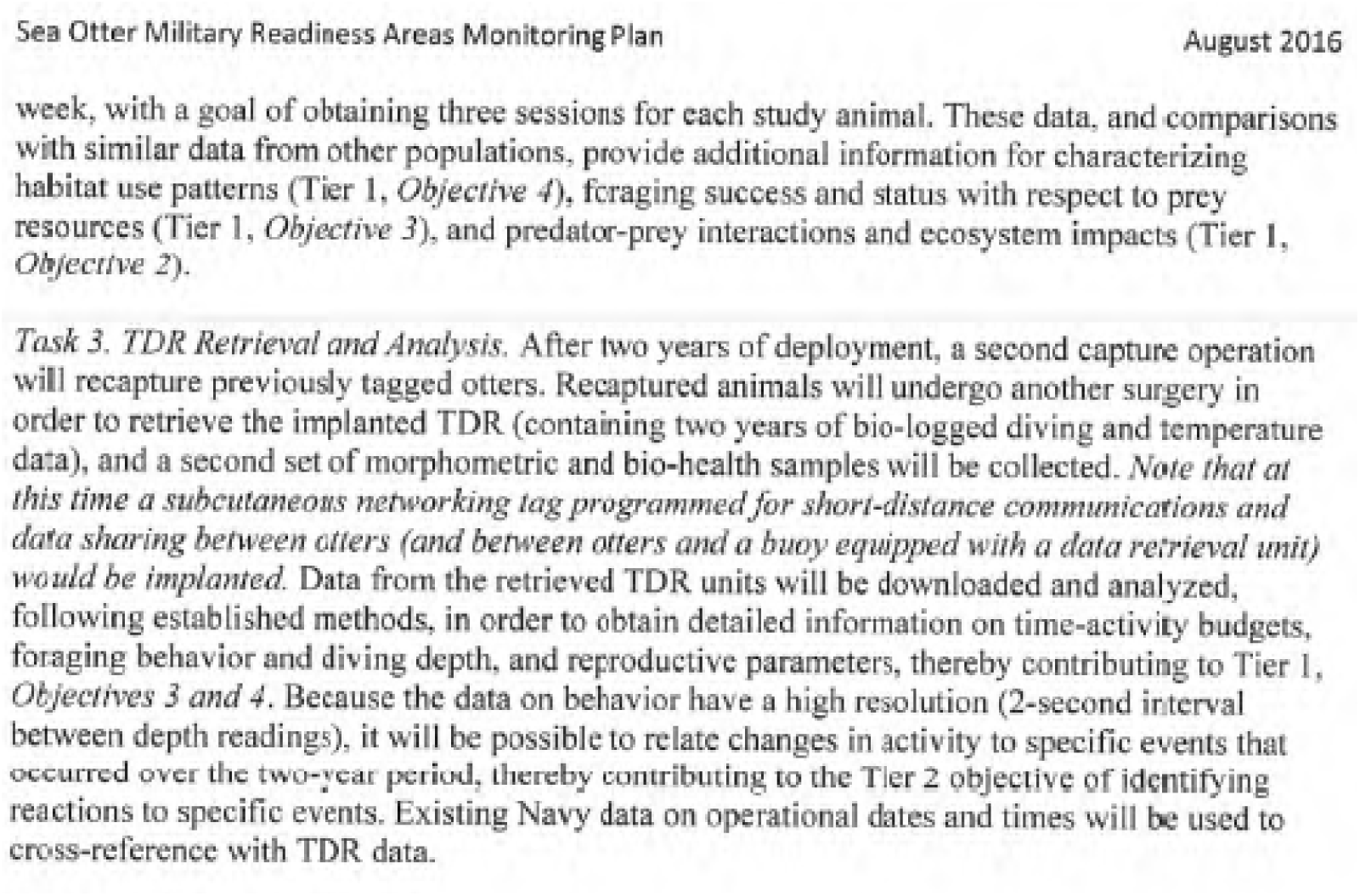
Page 9 of Monitoring and Research Plan for Southern Sea Otter Military Readiness Area.

**Figure 1.10.**
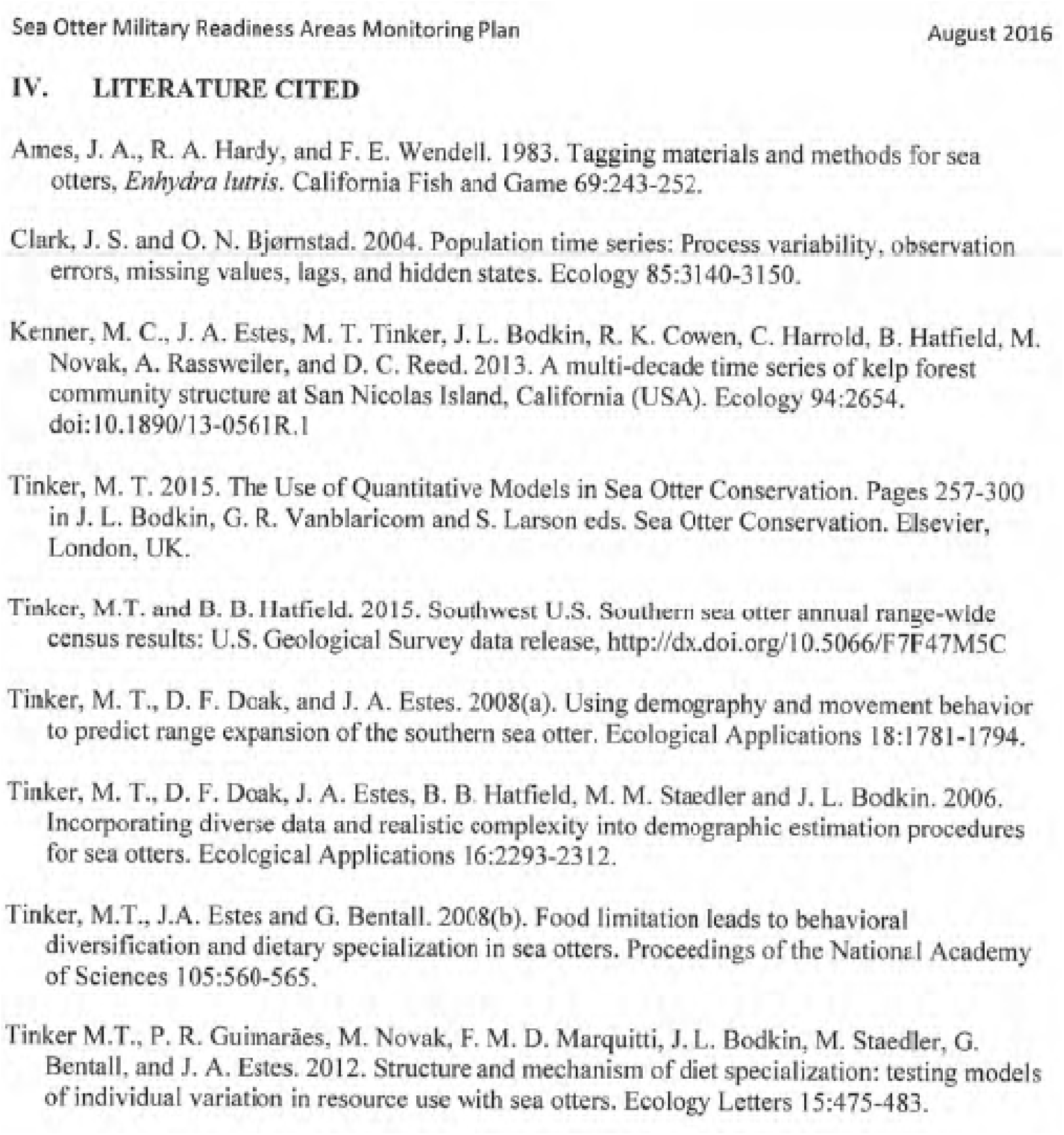
References for Monitoring and Research Plan for Southern Sea Otter Military Readiness Area.

**Figure 1.11.**
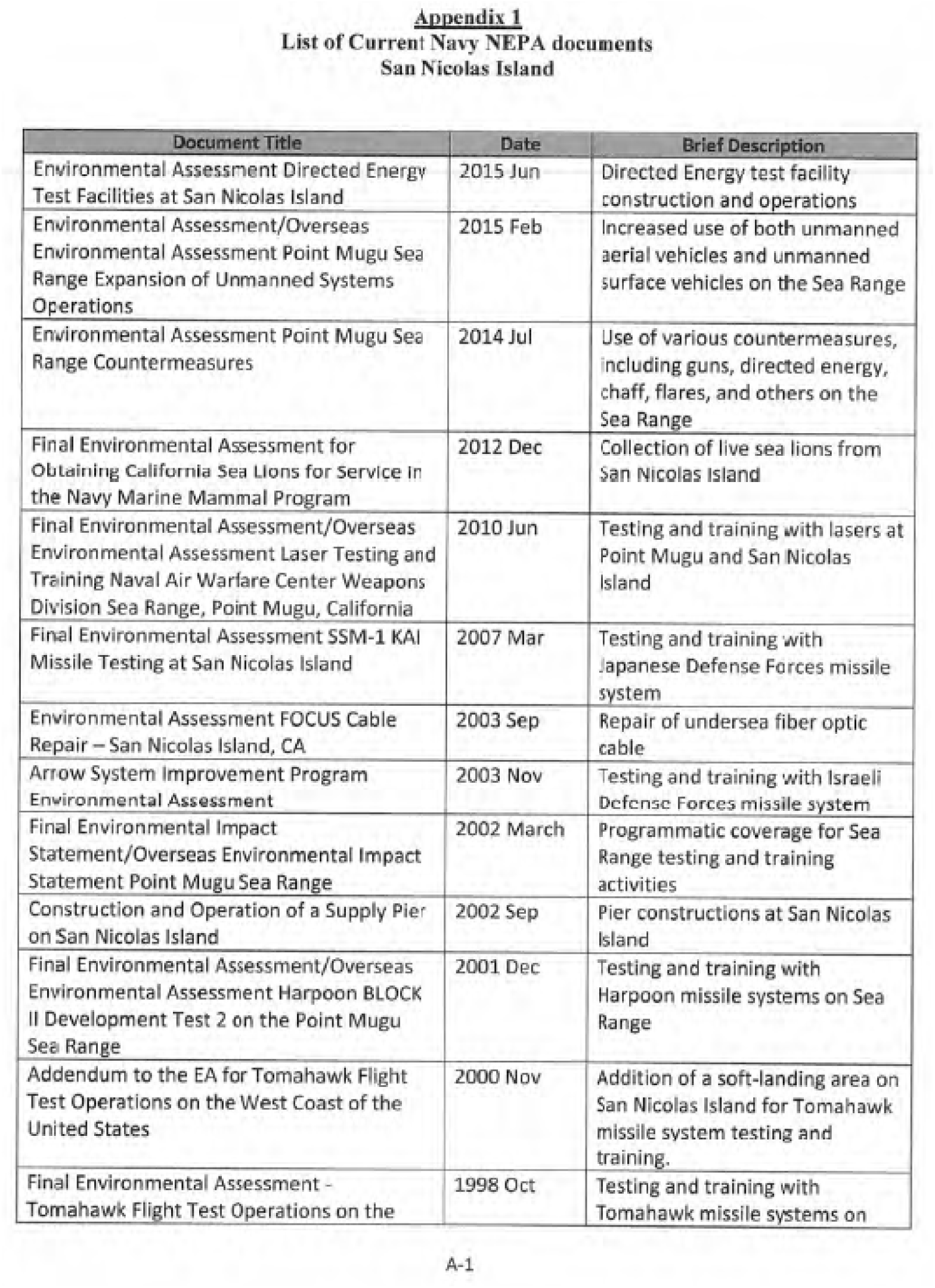
Page A-1 of appendix 1 for Monitoring and Research Plan for Southern Sea Otter Military Readiness Area.

**Figure 1.12.**
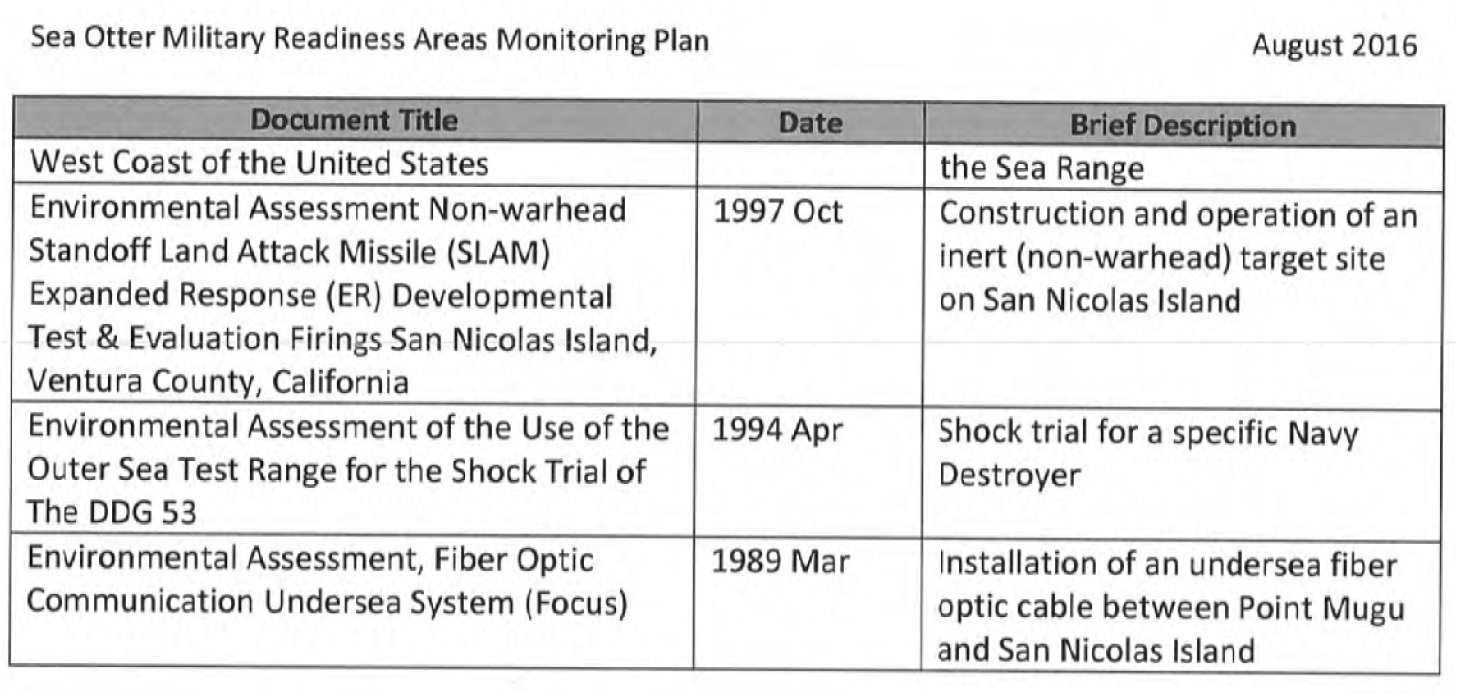
Page A-2 of appendix 1 for Monitoring and Research Plan for Southern Sea Otter Military Readiness Area.

## References Cited

Bodkin, J.L., 2015, Historic and contemporary status of sea otters in the North Pacific, chap. 3 of Larson, S.E., Bodkin, J.L., and VanBlaricom, G.R., eds., Sea otter conservation (1st ed.): London, Academic Press, p. 43–61, 10.1016/B978-0-12-801402-8.00003-2.

Estes, J.A., 1990, Growth and equilibrium in sea otter populations: Journal of Animal Ecology, v. 59, no. 2, p. 385–401, 10.2307/4870.

Estes, J.A., and Jameson, R.J., 1988, A double-survey estimate for sighting probability of sea otters in California: The Journal of Wildlife Management, v. 52, no. 1, p. 70–76, 10.2307/3801061.

Estes, J.A., Jameson, R.J., and Johnson A.M., 1981, Food selection and some foraging tactics of sea otters: Pages 606-641 in Chapman J.A. and Pursley D., editors, Worldwide Furbearer Conference Proceedings, Volume 1. University of Maryland Press, Baltimore, Maryland.

Estes, J.A., Riedman, M.L., Staedler, M.M., Tinker, M.T., and Lyon, B.E., 2003, Individual variation in prey selection by sea otters—Patterns, causes and implications: Journal of Animal Ecology, v. 72, no. 1, p. 144–155, 10.1046/j.1365-2656.2003.00690.

Hatfield, B.B., 2005, The translocation of sea otters to San Nicolas Island: an update, *in* Garcelon D.K. and Schwemm C.A., eds., Proceedings of the Sixth California Islands Symposium, Ventura, California, December 1–3, 2003, p. 473–475.

Hatfield, B.B., Yee J.L., Kenner, M.C., Tomoleoni J.A., 2019, California sea otter (Enhydra lutris nereis) census results, spring 2019: U.S. Geological Survey Data Series 1118, 12 p., 10.3133/ds1118.

Kenner, M.C., and Tomoleoni, J.A., 2020, Kelp forest monitoring at Naval Base Ventura County, San Nicolas Island, California—Fall 2018 and Spring 2019, fifth annual report: U.S. Geological Survey Open-File Report 2020–1091, 93 p., 10.3133/ofr20201091.

Kenner, M.C., and Tomoleoni, J.A., 2021, Kelp forest monitoring at Naval Base Ventura County, San Nicolas Island, California—Fall 2019, sixth annual report: U.S. Geological Survey Open-File Report 2021–1081, 111 p., 10.3133/ofr20211081.

McCullagh, P., and Nelder, J., 1989, Generalized linear models (Monographs on statistics and applied probability, 2d ed.): London, Chapman & Hall/CRC, 10.1201/9780203753736.

Newsome, S.D., Bentall, G.B., Tinker, M.T., Oftedal, O.T., Ralls, K., Estes, J.A., and Fogel, M.L., 2010, Variation in δ^13^C and δ^15^N diet–Vibrissae trophic discrimination factors in a wild population of California sea otters: Ecological Applications, v. 20, no. 6, p. 1744–1752, 10.1890/09-1502.1.

Oftedal, O.T., Ralls, K., Tinker, M.T., and Green, A., 2007, Nutritional constraints on the southern sea otter in the Monterey Bay National Marine Sanctuary, and a comparison to sea otter populations at San Nicolas Island, California and Glacier Bay, Alaska: Joint final report to the Monterey Bay National Marine Sanctuary and the Marine Mammal Commission, 23 p.

R Core Team, 2025, The R project for statistical computing. R Foundation for Statistical Computing, Vienna, Austria, https://www.R-project.org/

Ralls, K., Hatfield B.B., and Siniff D.B., 1995, Foraging patterns of California sea otters as indicated by telemetry: Canadian Journal of Zoology 73:523-531, 10.1139/z95-060.

Rathbun, G.B., Hatfield, B.B., and Murphey, T.G., 2000, Status of translocated sea otters at San Nicolas Island, California: The Southwestern Naturalist, v. 45, no. 3, p. 322–328, 10.2307/3672835.

Tinker, M.T., 2015, The use of quantitative models in sea otter conservation, chap. 10 of Larson, S.E., Bodkin, J.L., and VanBlaricom, G.R., eds., Sea otter conservation (1st ed.): London, Academic Press, p. 257–300, 10.1016/B978-0-12-801402-8.00010-X.

Tinker, M.T., Estes, J.A., Ralls, K., Williams, T.M., Jessup, D., and Costa, D.P., 2006, Population dynamics and biology of the California sea otter (*Enhydra lutris nereis*) at the southern end of its range: Santa Barbara, California, University of California, Coastal Research Center, Marine Science Institute, Minerals management Service Pacific Outer Continental Shelf Region Study 2006–007, Minerals management Service Cooperative Agreement Number 14-35-0001-31063.

Tinker, M.T., Guimarães, P.R., Novak, M., Marquitti, F.M.D., Bodkin, J.L., Staedler, M., Bentall, G., and Estes, J.A., 2012, Structure and mechanism of diet specialisation—Testing models of individual variation in resource use with sea otters: Ecology Letters, v. 15, no. 5, p. 475–483, 10.1111/j.1461-0248.2012.01760.x.

Tinker, M.T., Bentall, G., and Estes, J.A., 2008, Food limitation leads to behavioral diversification and dietary specialization in sea otters: Proceedings of the National Academy of Sciences of the United States of America, v. 105, no. 2, p. 560–565, 10.1073/pnas.0709263105.

Tinker, M.T., Tomoleoni, J.A., Weitzman, B.P., Staedler, M., Jessup, D., Murray, M.J., Miller, M., Burgess, T., Bowen, L., Miles, A.K., Thometz, N., Tarjan, L., Golson, E., Batac, F., Dodd, E., Berberich, E., Kunz, J., Bentall, G., Fujii, J., Nicholson, T., Newsome, S., Melli, A., LaRoche, N., MacCormick, H., Johnson, A., Henkel, L., Kreuder-Johnson, C., and Conrad, P., 2019, Southern sea otter (*Enhydra lutris nereis*) population biology at Big Sur and Monterey, California—Investigating the consequences of resource abundance and anthropogenic stressors for sea otter recovery: U.S. Geological Survey Open-File Report 2019–1022, 225 p., 10.3133/ofr20191022.

U.S. Fish and Wildlife Service, 2012, Final supplemental environmental impact statement on the translocation of southern sea otters: Ventura, California, U.S. Fish and Wildlife Service, Ventura Fish and Wildlife Office, 348 p.

Wickham, H., 2016, ggplot2—Elegant graphics for data analysis (2d ed.): New York, Springer-Verlag, 212 p.

Yee, J.L, Tomoleoni, J.A., Kenner, M.C., Fujii, J., Bentall G.B., Tinker, M.T., and Hatfield, B.B., 2020, Southern (California) sea otter population status and trends at San Nicolas Island, 2017-2020: U.S. Geological Survey Open_File Report 2020-1115, 38p., accessed April 15, 2026, at 10.3133/ofr20201115.

Yee, J.L, Tomoleoni, J.A., Kenner, M.C., Fujii, J.A., Bentall, G.B., Staedler, M.M., and Hatfield, B.B., 2023, Southern (California) Sea Otter Population Status and Trends at San Nicolas Island, 2020-2023: U.S. Geological Survey Open File Report 2023-1071, 37p., accessed April 15, 2026, at 10.3133/ofr20231071.

